# The *Drosophila* Cortactin Binding Protein 2 homolog, Nausicaa, regulates lamellipodial actin dynamics in a Cortactin-dependent manner

**DOI:** 10.1101/376665

**Authors:** Meghan E. O’Connell, Divya Sridharan, Tristan Driscoll, Ipsita Krishnamurthy, Wick G. Perry, Derek A. Applewhite

## Abstract

*Drosophila* CG10915 is an uncharacterized protein coding gene with sequence similarity to human Cortactin Binding Protein 2 (CTTNBP2) and Cortactin Binding Protein 2 N-terminal like (CTTNBP2NL). We have named this gene *Nausicaa (naus)* and characterize it through a combination of quantitative live-cell total internal reflection fluorescence (TIRF) microscopy, electron microscopy, RNAi depletion, and genetics. We found that Naus co-localizes with F-actin and Cortactin in the lamellipodia of *Drosophila* S2R+ and D25c2 cells and this localization is lost following Cortactin or Arp2/3 depletion or by mutations that disrupt a conserved proline patch found in its mammalian homologs. Using Permeabilization Activated Reduction in Fluorescence (PARF) and Fluorescence Recovery after Photo-bleaching (FRAP), we find that depletion of Cortactin alters Naus dynamics leading to a decrease in its half-life. Furthermore, we discovered that Naus depletion in S2R+ cells led to a decrease in actin retrograde flow and a lamellipodia characterized by long, unbranched filaments. We demonstrate that these alterations to the dynamics and underlying actin architecture also affect D25c2 cell migration and decrease arborization in *Drosophila* neurons. We present the novel hypothesis that Naus functions to slow Cortactin’s disassociation from Arp2/3 nucleated branch junctions, thereby increasing both branch nucleation and junction stability.

## Introduction

Cell migration is critical to a number of physiological processes including wound healing and immune function, development, neurogenesis, and vascularization. Aberrant cell migration is also the cause of a number of diseases including schizophrenia and mental disabilities, immunodeficiency, craniofacial disorders, and metastasis (Ridley et al., 2003). Cell migration relies heavily on the actin cytoskeleton, and the regulation of actin dynamics has major consequences for the underlying actin architecture dictating how cells migrate. Migration proceeds in four major steps - protrusion, adhesion, contraction, and retraction (Ridley et al., 2003). During the protrusion step of cell migration, the cell generates two major types of actin-based structures: lamellipodia and filopodia. While filopodia are characterized by parallel, unbranched actin filaments (Svitkina et al., 2003), the lamellipodia is composed of a densely branched network of actin filaments forming a sheet-like exploratory organelle (Abercrombie, M. et al., 1970). One protein that defines the lamellipodia is the actin-related protein 2/3 (Arp2/3) complex which generates new branches from the sides of pre-existing filaments resulting in a highly branched actin network (Machesky et al., 1999; Mullins et al., 1997; Suraneni et al., 2012; Svitkina and Borisy, 1999). It is the addition of actin subunits (G-actin), spread across the entire expanse of the lamellipodia that leads to protrusion of this organelle. The Arp2/3 complex must be activated by proteins known as nucleation promoting factors (NPFs) in order to nucleate filaments (Machesky et al., 1999; Prehoda et al., 2000; Zalevsky et al., 2001). NPFs have been divided into two types: the WASP/N-WASP and the SCAR/WAVE family of proteins comprising type 1 NPFs, and Cortactin and the closely related hematopoetic-specific protein-1 (HS1) comprising type II (Goley and Welch, 2006). While type I NFPs generally bind and activate Arp2/3 via a shared VCA (verprolin homology, central, acidic) region, Cortactin and HS1 use an N-terminal acidic region (NtA) (Goley and Welch, 2006; Weaver et al., 2001).

Cortactin, unlike type I NPFs, can be found integrated within the lamellipodia. Data from FRAP analysis suggests that it recovers throughout the organelle after photobleaching rather than just at the leading edge (Lai et al., 2008). Cortactin can bind to both the sides of actin filaments and at Arp2/3 generated branch junctions where it is thought to stabilize them (Weaver et al., 2001). Interestingly, *in vitro* single molecule experiments determined that Cortactin has a ~300 fold increased affinity for branch junctions over the sides of actin filaments suggesting the protein preferentially targets these sites (Helgeson and Nolen, 2013). Type I NPFs are more potent activators of the Arp2/3 complex than Cortactin, however, the addition of Cortactin to GST-VCA beads increased bead motility, suggesting that Cortactin may synergize with type I NPFs during filament nucleation (Helgeson and Nolen, 2013; Siton et al., 2011; Weaver et al., 2002). Previously, it had been shown that Cortactin competes with the VCA domain for binding to the Arp3 subunit of the Arp2/3 complex, and more recently single molecule experiments from Helgeson and Nolen demonstrate that Cortactin replaces the VCA domain of type I NPFs during nucleation (Helgeson and Nolen, 2013; Weaver et al., 2001). Thus, it appears that Cortactin both stimulates the formation of branches while simultaneously stabilizing them. This type of synergy may allow for continued dendritic nucleation while preventing the potential stalls caused by the tight membrane association of type 1 NPFs (Helgeson and Nolen, 2013). An examination of this synergy between type I and type II NPFs remains to be fully investigated *in vivo*, thus it is unclear how it fits into the paradigm of lamellipodial protrusion and cell migration.

Overexpression of Cortactin has been associated with increased metastasis and invasion in a host of cancers from breast carcinomas, and head and neck squamous cell carcinomas, to melanoma, colorectal cancers, and glioblastomas (Åkervall et al., 1995; Buday and Downward, 2007; Hirakawa et al., 2009; Kirkbride et al., 2011; Rothschild et al., 2006; Weaver, 2008; Xu et al., 2010). In support of this, overexpression of Cortactin in NIH 3T3 cells led to an increase in motility and invasiveness. Similarly, overexpression of Cortactin in breast cancer cells led to increased metastasis in nude mice (Patel et al., 1998). RNAi experiments in HT1080 cells suggest that Cortactin enhances lamellipodial persistence, and both the Arp2/3 and F-actin binding sites of Cortactin were required for this persistence (Bryce et al., 2005). Cortactin depletion also led to a decrease in the rate of adhesion formation, however, given the importance of the lamellipodia to the formation of nascent adhesions, it may be difficult to uncouple these phenotypes (Bryce et al., 2005; Wu et al., 2012).

Cortactin also localizes to other parts of the cell where dynamic actin assembly occurs including endosomes, podosomes, invadopodia, and the dendritic spines of neurons (Ammer and Weed, 2008; Buday and Downward, 2007; MacGrath and Koleske, 2012; Ren et al., 2009). Coincident with Cortactin at some of these sites of dynamic actin are two Cortactin binding proteins, Cortactin Binding Protein 2 (CTTNBP2) and Cortactin Binding Protein N-terminal like (CTTNBP2NL or CortBP2NL).

Human CTTNBP2, coded for by the *CTTNBP2* gene, is found primarily in neurons. CTTNBP2 interacts with the C-terminal SH3 domain of Cortactin (Ohoka and Takai, 1998) and previous studies have demonstrated that CTTNBP2 co-localizes with both Cortactin and actin at the lamellipodia. CTTNBP2 depletion in rat hippocampal neurons decreased the width and density of dendritic spines, suggesting that CTTNBP2 plays a role alongside with Cortactin in dendritic spine maintenance (Chen and Hsueh, 2012). Additionally, before dendritic spine formation, CTTNBP2 associates with microtubules through its central region and oligomerizes through its N-terminal region coiled-coil motif. CTTNBP2 oligomers bound to microtubules promotes microtubule bundle formation and tubulin acetylation (Shih et al., 2014).

Much less is known about CTTNBP2NL, and a clear cellular function for the protein has yet to be fully elucidated. CTTNBP2NL is found in epithelial, spleen, and liver cells and unlike CTTNBP2, CTTNBP2NL does not associate at the cell cortex, but instead can be found on actin stress fibers where it can redistribute Cortactin to these structures (Chen et al., 2012). Interestingly, in rat hippocampal neurons, exogenous CTTNBP2NL is unable to rescue the effects of CTTNBP2 depletion on dendritic spine morphology (Chen et al., 2012), indicating that the two proteins are not functionally similar in the context of mammalian dendritic spine morphology. Given Cortactin’s widespread expression, CTTNBP2NL may very well play important roles in other dynamic actin-based structures in non-neuronal cell types.

Drosophila CG10915 is an uncharacterized protein-coding gene that shows amino acid sequence similarity to CTTNBP2 and CTTNBP2NL. CG10915 is expressed ubiquitously throughout the larval and adult fly, with higher expression levels in the central nervous system and ovaries (Gelbart and Emmert, 2013). Here, we investigate the role of CG10915 in *Drosophila* to determine its role in actin dynamics. We demonstrate that the *Drosophila* gene CG10915 (hereafter referred to as *Nausicaa* (*naus*)) alters lamellipodial and protrusive actin dynamics in migratory cells and neurons in a Cortactin-dependent manner.

## Results

Bioinformatic queries initially indicated that the *Drosophila* CG10915 locus at cytological position 55B9, was a potential homolog of human Filamin-A interacting protein (FILIP) due to it sharing approximately 20 percent identity. Upon further refinement of these queries (Clustal Omega Multiple Sequence Alignment, Goujon et al., 2010; Sievers et al., 2011) we found that CG10915 is more similar to both human Cortactin Binding Protein 2 N-terminal like (CTTNBP2NL) and Cortactin Binding Protein 2 (CTTNBP2) sharing approximately 30 percent and 28 percent identity, respectively (Supplemental Figure 1). We have subsequently named CG10915, the putative *Drosophila* homolog of CTTNBP2 and CTTNBP2NL, *nausicaa* (*naus*), after the princess in Homer’s *The Odyssey* who helps to ensure Odysseus’s safe passage home from Phaeacia. *Naus* has two splice variants, each encoding a polypeptide of 609 amino acids. The highest degree of conservation between Naus, CTTNBP2, and CTTNBP2NL occurs in the coiled-coil motif found within the N-terminal Cortactin binding protein (CortBP2) domain. Furthermore, the three proteins also share a highly conserved proline-rich patch located near their C-termini that has been shown to facilitate the interaction between Cortactin and CTTNBP2 in COS cells (Supplemental Figure 1) (Chen et al., 2012).

### Nausicaa localizes to lamellipodia of S2R+ cells in a Cortactin-dependent manner

We first assessed the localization of EGFP-tagged Naus by live-cell imaging of *Drosophila* S2R+ cells using total internal reflection fluorescence (TIRF) microscopy (Figure 1A, Supplemental Video 1). Interestingly, we observed an enrichment of Naus in the circumferential lamellipodia of these cells which persisted following fixation (data not shown) and when we extracted the cells with detergent prior to fixation (Figure 1B). We next investigated whether Naus localized to other actin-based structures. *Drosophila* S2R+ cells do not form prominent stress fibers, however, ML-DmD25c2 (D25) cells which are derived from third instar imaginal wing discs, readily form these structures. We co-expressed EGFP-tagged Naus with mCherry-Alpha-actinin to mark stress fibers and actin bundles, and observed Naus weakly localizing to these structures as well (Supplemental Figure 2A). Furthermore, much like our results from S2R+ cells, we also observed lamellipodial enrichment of Naus in D25 cells (Supplemental Figure 2B). Interestingly, while CTTNBP2NL can also be found co-localizing to microtubules in COS cells (Chen et al., 2012), we failed to observe any colocalization between Naus and microtubules in either S2R+ or D25 cells under these conditions (data not shown). Collectively, these results suggest that Naus behaves similarly to both CTTNBP2 and CTTNBP2NL, localizing to both the lamellipodia and bundled actin structures.

**Figure 1.**
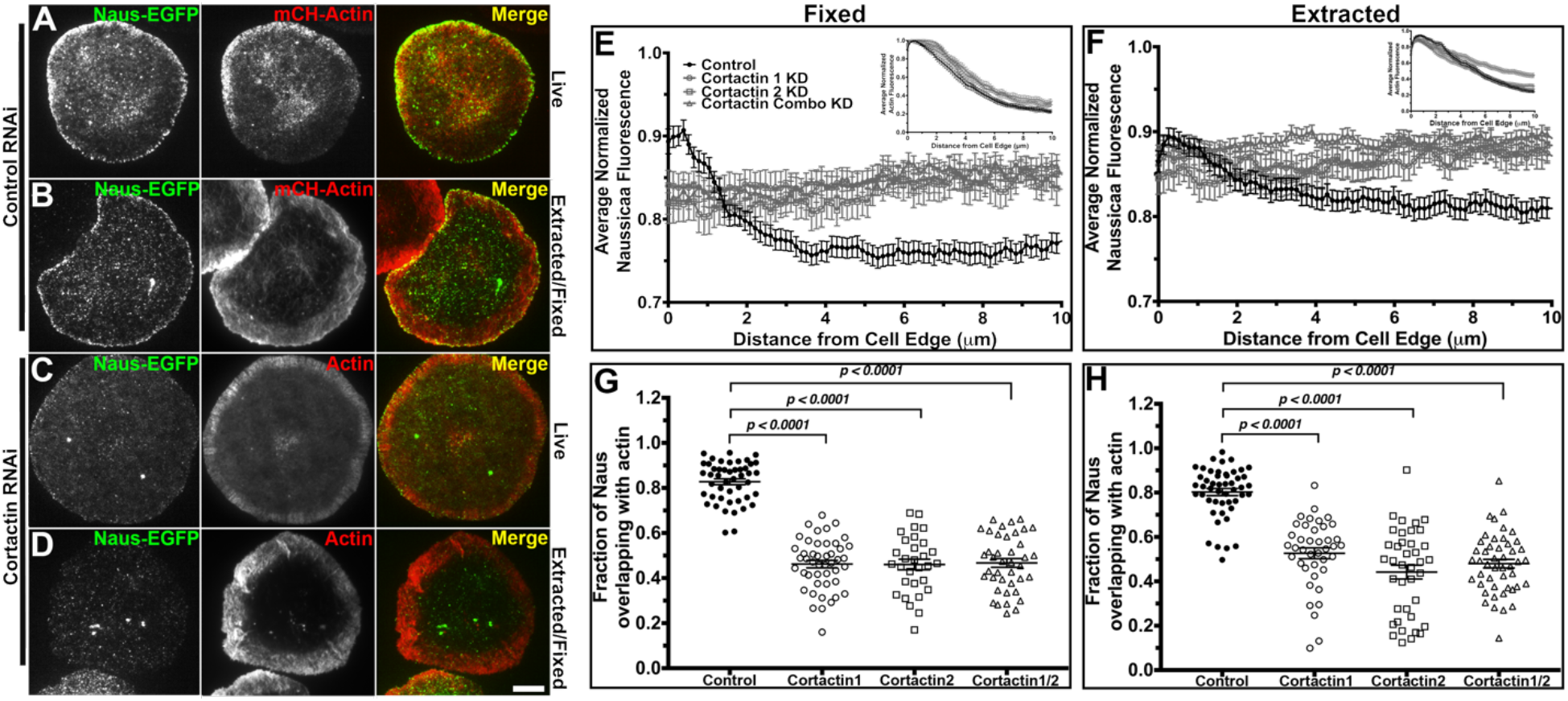
Nausicaa’s lamellipodial localization is Cortactin-dependent. (A & B) Control RNAi treated S2R+ cells transfected with Naus-EGFP (left panels, green in merged image) and mCherry-Actin (middle panels, red in the merged image) imaged live (A) or extracted with detergent prior to fixation and stained with phalloidin (middle panels, red in the merged image) to visualize F-actin (B). (C & D) Cortactin RNAi treated S2R+ cells transfected with Naus-EGFP (left panels, green in the merged images) and mCherry-Actin (middle panels, red in the merged image) imaged live (C) or extracted with detergent prior to fixation and stained with phalloidin (middle panels, red in the merged images) to visualize F-actin (D). Scale bar = 10 μm. (E) Line scan analysis of the lamellipodial distribution of Naus-EGFP in fixed S2R+ cells treated with control RNAi (black circles), or Cortactin RNAi from two independent RNAi targets (open gray circles or squares) or the combination of the two targets (open gray triangles). Inset is the corresponding averaged normalized actin fluorescence. Error bars denote S.E.M. (F) Line scan analysis of the lamellipodial distribution of Naus-EGFP in cells that were extracted with detergent prior to fixation following control RNAi (black circles) or treatment with two independent Cortactin RNAi targets (open gray circles or squares) or the combination of the two (open gray triangles). Inset is the corresponding averaged normalized actin fluorescence. Error bars denote S.E.M. (G) Mander’s coefficient of the amount of overlap between Naus-EGFP and F-actin stained by phalloidin. Cells were treated with control RNAi (black circles) or one of two Cortactin RNAi targets (open gray circles or squares) or the combination of the two (open gray triangles). There was a statistically significant decrease in lamellipodial enrichment when cells were depleted of Cortactin (error bars = S.E.M., n = 30-45 cells per condition) (*** = p-value <0.0001, Student’s t-test). (H) Mander’s coefficient quantifying the amount of overlap between Naus-EGFP and F-actin as visualized by phalloidin in cells that were first extracted prior to fixation. control RNAi is shown in black circles, Cortactin RNAi (from two independent targets) is shown in open gray circles or squares, and the combination of the two Cortactin RNAi targets is shown in open gray triangles. There was a statistically significant decrease in the amount of overlap between Naus-EGFP and F-actin stained by phalloidin (error bars are S.E.M, n= 40-50 cells per condition) (*** = p-value < 0.0001, Student’s t-test).

Given the potential interaction between Naus and *Drosophila* Cortactin (CG3637), we next tested whether this lamellipodial enrichment in S2R+ cells is Cortactin-dependent. While it has been demonstrated that mammalian CTTNBP2 and CTTNBP2NL interacts with Cortactin, the role Cortactin plays in this interaction is unclear. Using two independent dsRNA sequences we depleted Cortactin and expressed EGFP-tagged Naus and observed a distinct loss on Naus’ lamellipodial localization (Figure 1C, and Supplemental Video 1). This loss in enrichment was even more evident in cells that were detergent extracted prior to fixation (Figure 1D). Line-scan analysis, where we compared control RNAi treated cells to cell treated with either Cortactin dsRNAs or in combination, further corroborated this change in localization (Figure 1E). To quantify this change, we used Mander’s Coefficient and measured the fraction of Naus overlapping with F-actin (stained by fluorescently labeled phalloidin) following Cortactin depletion, and found a statistically significant decrease in the amount of Naus overlapping with actin further supporting that Naus’ association with actin cytoskeleton is Cortactin-dependent (Bolte and Cordelières, 2006) (Figure 1G & H). This differs from CTTNBP2 where upon Cortactin re-distribution, CTTNBP2 does not re-localize in neurons suggesting a Cortactin-independent mechanism of localization for this potential Naus homolog (Chen and Hseuh, 2012). Given that Cortactin interacts with Arp2/3 complex at the lamellipodia (Uruno et al., 2001), we depleted the p20 subunit of Arp2/3 complex by RNAi and observed a similar loss of localization (Supplemental Figure 3). Collectively, these results suggest that Naus is enriched in the lamellipodia and that this enrichment to actin structures is Cortactin-dependent.

Given that Naus’ lamellipodial localization is Cortactin-dependent, we next wanted to characterize the relationship between the two proteins. We co-expressed Naus-EGFP with myc-tagged Cortactin in S2R+ cells and again used Mander’s Coefficient to determine the degree of overlap between these proteins (Figure 2A, B, & D). The Mander’s Coefficient revealed that just over 50% of myc-Cortactin overlapped with Naus while nearly 80% of Naus-EGFP overlapped with Cortactin. This asymmetry in co-localization, which was statistically significant (Student’s t-test, p-value <0.0001), suggests that while not all of the Cortactin in the cell is associated with Naus, the majority of the Naus in the cell can be found overlapping with Cortactin. This supports the hypothesis that Naus relies on Cortactin for proper localization. Naus, like CTTNBP2 and CTTNBP2NL, has a proline rich patch (PPPIP) that was previously shown to be required for Cortactin binding (Supplemental Figure 1) (Chen et al., 2012). To further elucidate the relationship between Naus and *Drosophila* Cortactin we mutated these proline residues (amino acid positions 563-567) to alanine and expressed an EGFP-tagged version (Naus-AAAIA) in S2R+ cells (Figure 2C). Our initial observations indicated that rather than a specific localization to actin-based structures, Naus-AAAIA appeared to be distributed non-specifically throughout the cell a pattern similar to what we observe when we expressed untagged-EGFP in these cells (Figure 2C & E). Line-scan analysis corroborates this observation and reveals a distinct loss in lamellipodial-enriched Naus when these residues are mutated (Figure 2F). This loss is similar to the loss of lamellipodial enrichment we observed following Cortactin RNAi (Figure 1C). When we quantified the amount of colocalization by Mander’s Coefficient we observed a statistically significant decrease in the amount of overlap between Naus-AAAIA and Cortactin further supporting the observation that this proline patch is facilitating the interaction between Naus and Cortactin (Figure 2B & D). Similar to what we observed in S2R+ cells, EGFP-tagged Naus-AAAIA failed to localize specifically to actin structures in D25 cells (Supplemental Figure 2C). These results suggest that Naus, like its mammalian counterparts, interacts with Cortactin through this conserved proline patch, but uniquely, requires Cortactin for proper localization. Interestingly, while we failed to observe microtubule localization in cells expressing wild-type EGFP-Naus, on occasion we did observe EGFP-tagged Naus-AAAIA colocalizing with microtubules in both S2R+ and D25 cells (Supplemental Figure 2D). It is likely that under conditions where its affinity for Cortactin is reduced, Naus may bind microtubules. While more detailed analysis of this microtubule localization is needed, we feel that this is beyond the scope of this current study.

**Figure 2.**
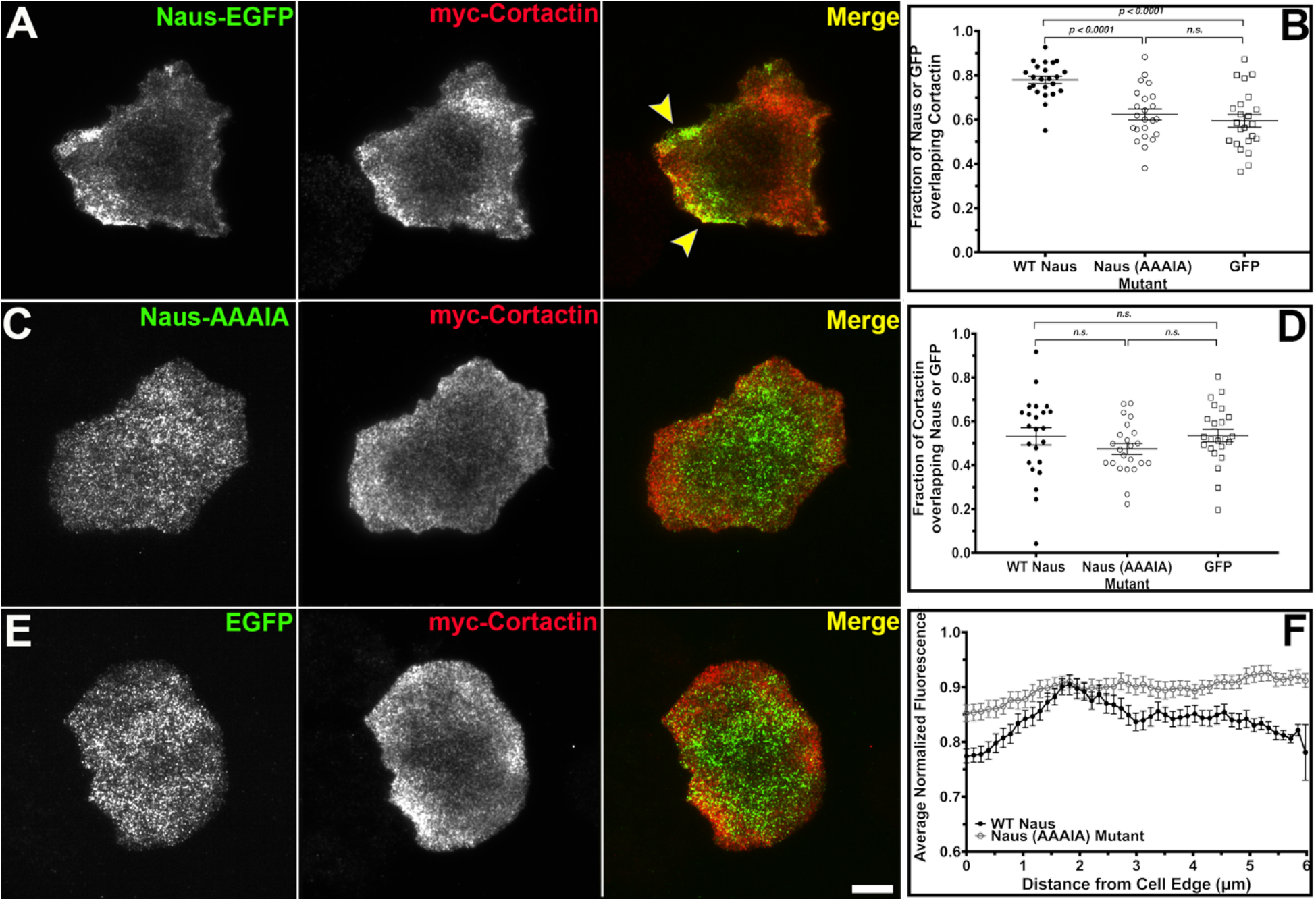
Nausicaa colocalizes with Cortactin through a conserved proline rich motif. (A-C) Fixed S2R+ cells co-transfected with myc-Cortactin and either Naus-EGFP (A), Naus(AAAIA)-EGFP mutant (B), or EGFP control (C) (left panels, green in merged images) and stained with anti-Myc (middle panels, red in merged images). Yellow arrowheads in A indicate regions of colocalization between Naus-EGFP and myc-Cortactin. Scale bar = 10 μm. (D) Mander’s coefficient analysis for Naus or EGFP constructs overlapping with Myc-Cortactin (*** = p-value < 0.0001, One-way ANOVA, error bars = S.E.M., n=23 cells per condition). (E) Mander’s coefficient analysis for Cortactin overlapping with Naus or EGFP constructs. None of these differences were statistically significant (n.s.). (F) Line scan analysis of the lamellipodial distribution of WT Naus-EGFP (black circles) or Naus-EGFP AAAIA (open gray circles) in fixed S2R+ cells (error bars denote S.E.M., n= 23 cells per condition).

Given this dependence on Cortactin for its lamellipodial localization, we next sought to determine if Naus’ dynamics are altered in the absence of Cortactin. We first used Permeabilization Activated Reduction in Fluorescence (PARF) to measure the loss of Naus-EGFP fluorescence following control or Cortactin RNAi treatments (Figure 3). PARF uses a low concentration of digitonin to gently permeabilize cells which leads to a large-scale dilution of the unbound pool of protein and a disruption of the initial equilibrium of the bound protein. The subsequent decrease in fluorescence can be fit to an exponential model and can be used to calculate a t_1/2_ for fluorescence loss of the bound fraction (Singh et al., 2016). Cortactin depletion led to a rapid loss in Naus-EGFP fluorescence following permeabilization with an average half-time of 5.956 ± 0.8s while control RNAi treated cells had an average half-time of fluorescence decay of 11.28 ± 2.1s, nearly double that of Cortactin depleted cells (Figure 3A-B & E-F, Supplemental Videos 2-3). These results suggest that depletion of Cortactin leaves a larger portion of the Naus pool free to quickly diffuse out of the cell rather than maintaining an association with Cortactin and the actin cytoskeleton. To corroborate our PARF results, we also performed Fluorescence Recovery After Photobleaching (FRAP) in cells treated with control or Cortactin RNAi. Again, we found that depletion of Cortactin led to a decrease in the half-time of recovery, from 56.5 ± 12.2s in control cells to 24.7 ± 4.9s in Cortactin depleted cells (Figure 3C-D & G-H, Video 4). Similar to our PARF results, our FRAP experiments suggest that in the presence of Cortactin, Naus is more stably associated with the cytoskeleton leading to a slower half-time of recovery as compared to Cortactin depleted cells. Collectively, these results indicate that Cortactin may function as an anchor, helping Naus maintain lamellipodial localization.

**Figure 3.**
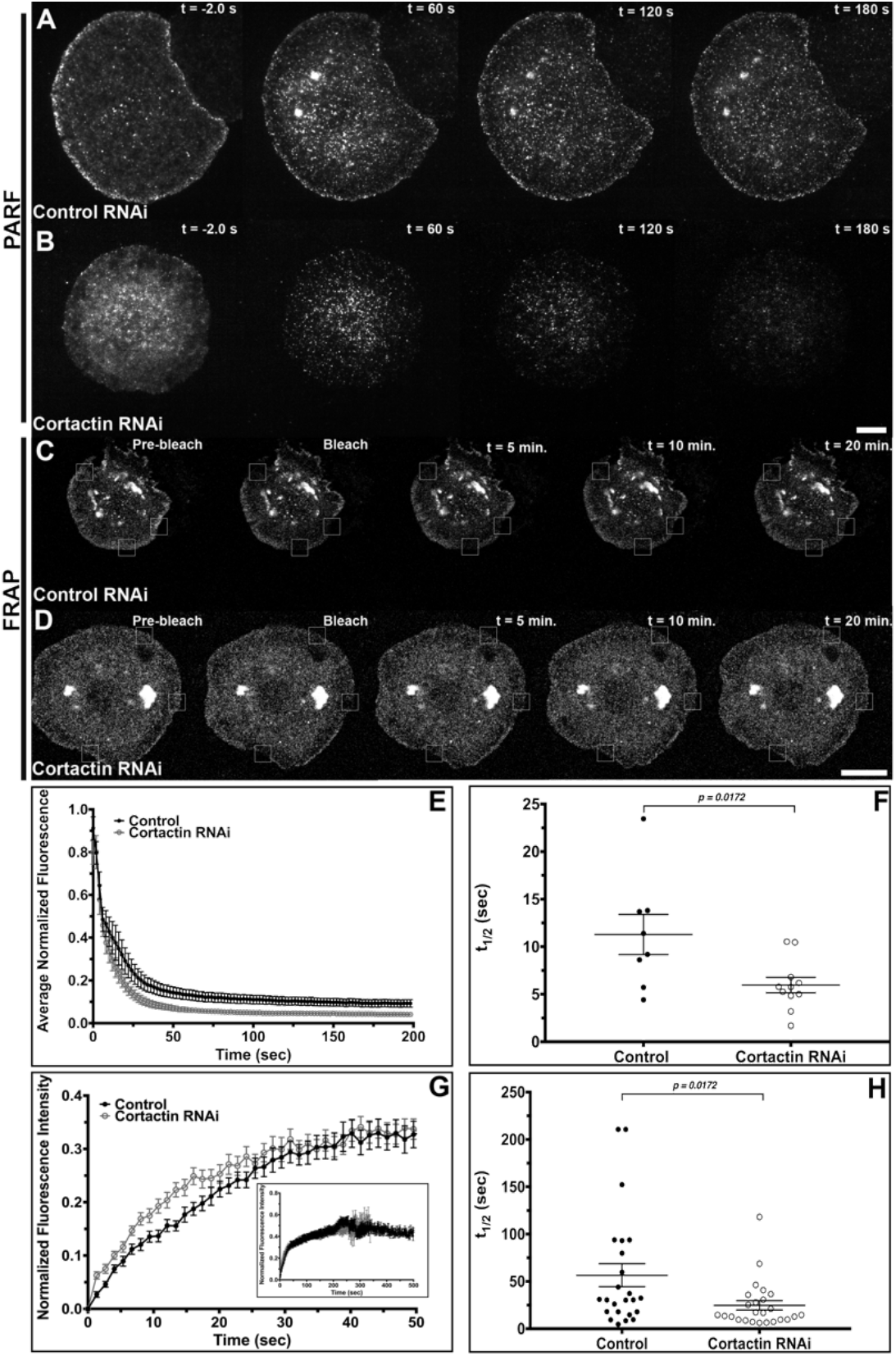
Depletion of Cortactin alters Nausicaa dynamics in the lamellipodia of S2R+ cells. (A-B) Time-lapse images of permeabilization activated reduction in fluorescence (PARF) in S2R+ cells transfected with Naus-EGFP following Control RNAi (A) or Cortactin RNAi (B) treatments. (C-D) Time-lapse images of fluorescence recovery after photobleaching (FRAP) of S2R+ cells transfected with Naus-EGFP following (C) Control RNAi or (D) Cortactin RNAi treatments. Small white boxes denote regions bleached. Scale bar = 10 μm (E) Average normalized fluorescence decay from PARF experiments. Error bars denote S.E.M. (F) Average half-life of Naus-EGFP fluorescence for Control (black circles) or Cortactin RNAi (open gray circles) conditions from PARF experiments. The fluorescent decay from Cortactin depleted cells was statistically significantly faster than control RNAi treated cells (p-value = 0.0172, Student’s t-test, Control: n = 8 cells, Cortactin RNAi: n = 11 cells). (G) Average normalized fluorescence recovery from FRAP experiments, in black circles, control RNAi and open gray circles, Cortactin RNAi. (H) Average half-life of Naus-EGFP fluorescence recovery for control (black circles) or Cortactin RNAi (open gray circles) treated cells from the FRAP experiments. Cortactin depleted cells recovered statistically significantly faster than control cells (p-value = 0.0178, Student’s t-test, Control: n = 24 cells, Cortactin RNAi: n = 25 cells). Error bars are S.E.M.

While our results indicate that Cortactin affects Naus’ dynamics, we wanted to determine if the inverse is also true. Once again, we used PARF, this time to measure the loss of Cortactin fluorescence following RNAi depletion of Naus (Figure 4A & B). This analysis revealed that depletion of Naus led to a statistically significant decrease in the half-time Cortactin’s fluorescence decay as compared to control RNAi treated samples (Figure 4C & D, Supplemental Videos 5-6). The average half-time of fluorescence decay for mCherry-Cortactin following Naus depletion was 15.32 ± 1.3 s, while the fluorescent decay in control treated cells was 20.13 ± 1.6 s (Figure 4C). A similar increase in Cortactin’s mobility was found in the dendritic spines of rat primary hippocampal neurons depleted of CTTNBP2 following FRAP analysis (Chen et al., 2012).

**Figure 4.**
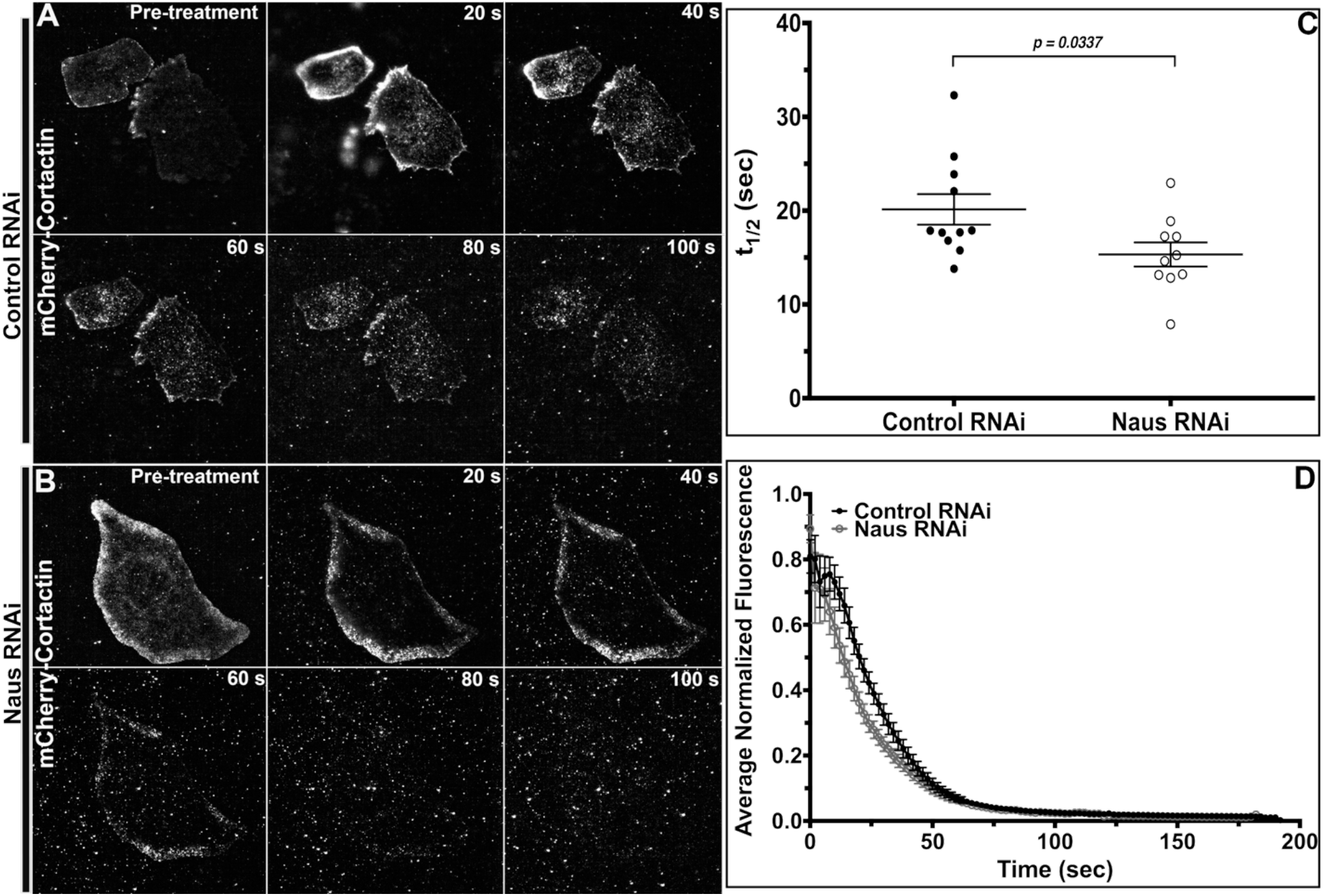
Depletion of Nausicaa alters Cortactin dynamics in S2R+ cells. (A & B) Time-lapse images of permeabilization activated reduction in fluorescence (PARF) of S2R+ cells transfected with mCherry-Cortactin following control RNAi (A) or Naus RNAi (B) treatments. (C) Average half-life of mCherry-Cortactin fluorescence for control (black circles) or Naus RNAi (open gray circles) treatments. Cortactin’s fluorescence decay was a statistically significantly faster following Naus RNAi as compared to control RNAi treated cells (*p-value 0.0337, Student’s t-test, Control: n = 10 cells, Naus RNAi: n = 11 cells). (D) Average normalized mCherry-Cortactin fluorescence decay from PARF experiments. Error bars denote S.E.M. Scale bar = 10 μm.

### Depletion of Nausicaa alters lamellipodial actin dynamics

Cortactin can function as both a type II nucleating promoting factor (NPF) and a stabilizer of Arp2/3 generated actin branches. Any changes to its dynamics could affect actin polymerization, branch density, and the overall rates of lamellipodial protrusion (Ammer and Weed, 2008; Bryce et al., 2005; Uruno et al., 2003; Weaver et al., 2001). Additionally, mammalian CTTNBP2 is known to regulate dendritic spine formation, which are Arp2/3-nucleated actin-rich structures (Chen et al. 2012), though the direct effect on actin dynamics and architecture is unexplored. Given the putative role Naus plays in regulating Cortactin dynamics, we sought to determine if Naus plays a role in regulating lamellipodial actin dynamics. We first depleted Naus in S2R+ cells and examined the circumferential lamellipodia of these cells by quantitative fluorescence microscopy (Figure 5A-D). Using phalloidin to measure F-actin we performed line-scan analysis as well as quantified the mean actin density of the lamellipodia (Figure 5C & D). Our analysis revealed an increase in actin fluorescence in the lamellipodia following Naus depletion which was statistically significant (p-value < 0.001, Student’s t-test, n= 30 cells per condition) when compared to control RNAi treated cells prepared in parallel (Figure 5D).

**Figure 5.**
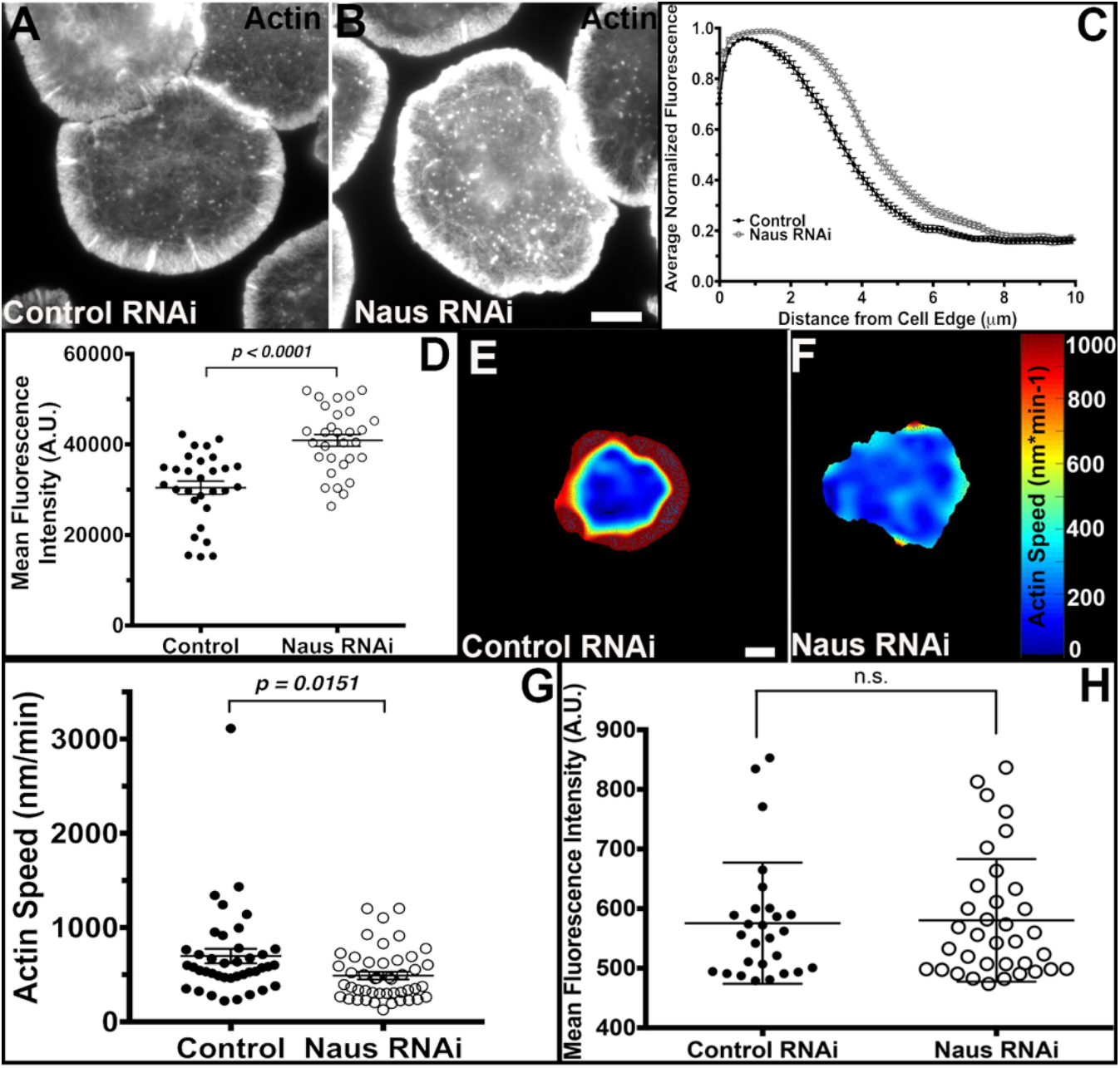
Nausicaa regulates lamellipodial actin density and actin retrograde flow in S2R+ cells. (A & B) Fixed S2R+ cells treated with control (A) or Naus (B) RNAi stained for F-actin with phalloidin. Gray levels have been set equal for comparison. Scale bars = 10 μm. (C) Line scan analysis of F-actin fluorescence in the lamellipodia from cells as shown in A & B. Fluorescence was normalized for each cell and averaged for each condition (black circles control RNAi and open gray circles Naus RNAi). (D) Mean fluorescence intensity of lamellipodial actin of cells treated with control RNAi (black circles) or Naus RNAi (open gray circles) from cells as shown in A & B. Naus RNAi led to a statistically significant increase in the F-actin fluorescence throughout the lamellipodial (**p-value <0.0001, Student’s t-test, n = 30 cells per condition). (E & F) Representative heat maps of actin speeds in control (E) or Naus (F) RNAi treated cells from QFSM analysis. Cool colors indicate slower rates of retrograde flow and warm colors represent faster speeds of actin retrograde flow. Scale bar = 10 μm. (G) Quantification of lamellipodial actin speeds from QFSM analysis. Naus depletion led to a statistically significant decrease in the rate of actin retrograde flow rates (*p-value 0.0151, Student’s t-test, Control RNAi: n = 40 cells, Naus RNAi: n = 46 cells). (H) Quantification of the mean fluorescence intensity of EGFP-Actin in the cells analyzed by QFSM (black circles control RNAi and open gray circles Naus RNAi). There was not a statistically significant difference between control and Naus RNAi treated cells.

As this increase in filamentous actin likely implies a change in dynamics, we next asked if depletion of Naus leads to changes in the rates of actin retrograde flow. Actin retrograde flow in the lamellipodia is the result of a combination of the plasma membrane pushing back against the force of actin polymerization and non-muscle myosin II contractility. Given Naus’ lamellipodial localization and its association with Cortactin, it is likely that any changes we observe in actin retrograde flow are due to changes in actin polymerization rather than contractility. To measure actin retrograde flow we turned to Quantitative Fluorescence Speckle Microscopy (QFSM) (Danuser and Waterman-Storer, 2006). Following treatment with Naus or control RNAi, we transfected S2R+ cells with EGFP-tagged actin using a copper-inducible promoter which allowed us to closely regulate the level of expression (Iwasa and Mullins, 2007). We imaged the cells by TIRF microscopy and analyzed the resulting movies using a Matlab-based program, QFSM, developed by the Danuser lab (Figure 5E & F, Supplemental Video 7) (Mendoza et al., 2012). Interestingly, Naus depletion led to a statistically significant 1.4-fold (p-value = 0.0151, Student’s t-test, n = 40 cells and 46 cell for control and Naus RNAi, respectively) decrease in actin retrograde flow speeds in the lamellipodia as compared to control RNAi treated cells (Figure 5G). We measured the mean fluorescence intensity of EGFP-actin in these cells to determine if actin expression levels were dictating the speed of retrograde flow and found no statistically significant difference between the two RNAi conditions (Figure 5H). This slowing of actin dynamics in combination with an increase in F-actin in the lamellipodia indicates that Naus helps to regulate actin branch dynamics, likely through its interaction with Cortactin.

### Depletion of Nausicaa leads to an increase in the number of long, unbranched actin filaments the lamellipodia

The decrease in the rate of actin polymerization coupled with the increase in filamentous actin we observed in the lamellipodia of Naus depleted S2R+ cells suggests that Naus may play a role in the regulating the fundamental architecture the actin cytoskeleton. To test this, we turned to platinum replica electron microscopy (Svitkina, T., 2016). We generated platinum replicas of control and Naus depleted S2R+ cells and imaged the actin cytoskeleton of the lamellipodia using electron microscopy (Figure 6). Interestingly, while the lamellipodia of control treated cells remained highly branched, typical of an Arp2/3 nucleated dendritic network (Figure 6A & B), the lamellipodia of Naus depleted cells was composed of extremely long, unbranched filaments with very few branch junctions (Figure 6C & D). Curiously, the ultrastructure of Naus depleted lamellipodia were reminiscent of the lamellipodia of Rat2 cells following the membrane targeting of Ena/VASP proteins (Bear et al. 2002), and suggest that in the absence of Naus there is either reduction in barbed-end capping or branching. Given that Cortactin is both an NPF and can stabilize the Arp2/3 complex at branch junctions, these long, unbranched filaments we observed are likely the result of a reduction in Cortactin’s activity. Without the stabilization provided by Naus, Cortactin fails to remain associated with Arp2/3 branches ultimately leading to reductions in actin polymerization and branch formation. Furthermore, these data assert a novel role for Naus in the regulation of the lamellipodial machinery and likely has broader implications in the way this actin-based protrusive organelle functions during cell migration.

**Figure 6.**
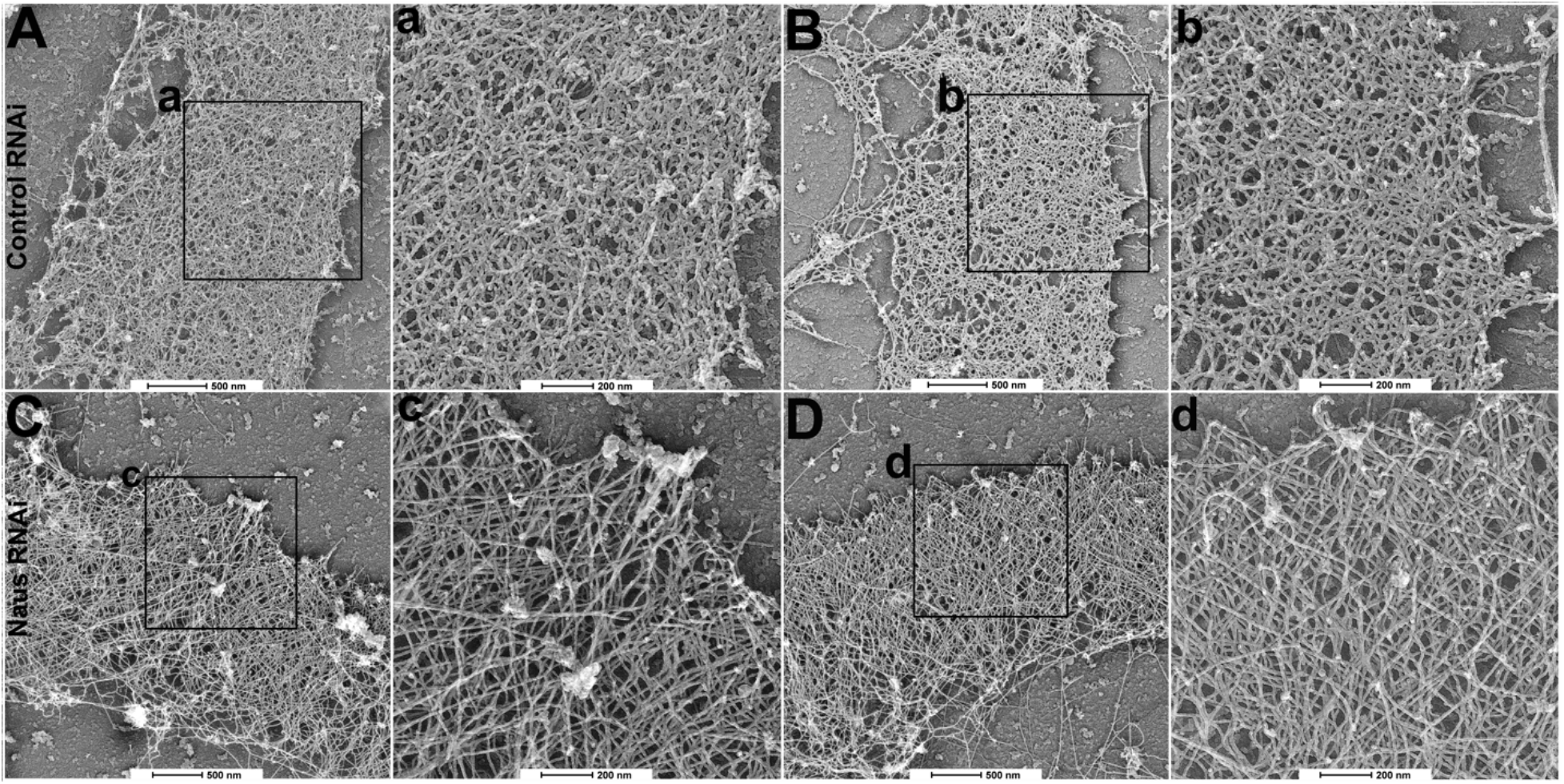
Depletion of Naus leads to an increase in the number of long, unbranched actin filaments on the lamellipodia. Platinum replicas of the lamellipodia of S2R+ cells treated with control (A & B) or Naus (C & D) RNAi. The black box denotes the region shown at higher magnification (control RNAi shown in a & b and Naus RNAi shown in c & d). Scale bars are given for each image. Scale bar in lower magnification images (A, B, C, D) 500 nm, scale bar in higher magnification images (a, b, c, d) 200 nm.

### Depletion of Nausicca leads to a decrease in cell migration and directionality

Actin polymerization is the main engine behind lamellipodial protrusion and ultimately, cell migration. Given the alterations to both actin dynamics and architecture we observed following Naus depletion, we next sought to determine whether Naus plays a role in cell migration. To do this, we performed a random cell migration assay where we treated D25 cells with control or Naus RNAi for seven days, plated them on mixture extracellular matrix (ECM), and imaged them by phase-contrast microscopy for six hours (Figure 7A-D, Supplemental Video 8). Cells were then manually tracked yielding migration speeds. In comparing the instantaneous velocity of approximately 50 cells per condition we found that depletion of Naus led to a modest but statistically significant decrease in the speed of cell migration. Control RNAi cells migrated at an average rate of approximately 1.6 ± 0.09 μm min^−1^ while Naus depleted cells migrated at an average rate of 1.4 ± 0.05 μm min^−1^(p-value < 0.0001, Student’s t-test) (Figure 7E). Cells undergoing random migration still maintain a degree of directionality. When we measured the directionality of Naus depleted cells we found they were statistically significantly less directional than Control RNAi treated cells. Where a value of 1.0 is completely directional, Control RNAi treated cells showed a value of 0.7 ± 0.07 a.u. while Naus depleted cells were 0.5 ± 0.04 a.u. (p= 0.0230, Student’s t-test) (Figure 7F). Directional persistence is a function of actin branch density and increases to actin branching positively correlates with the directionality of randomly migrating cells (Harms et al., 2005). Thus, the decrease in directionality we observed in Naus depleted D25 cells is consistent with the ultrastructural data gathered from S2R+ cells.

**Figure 7.**
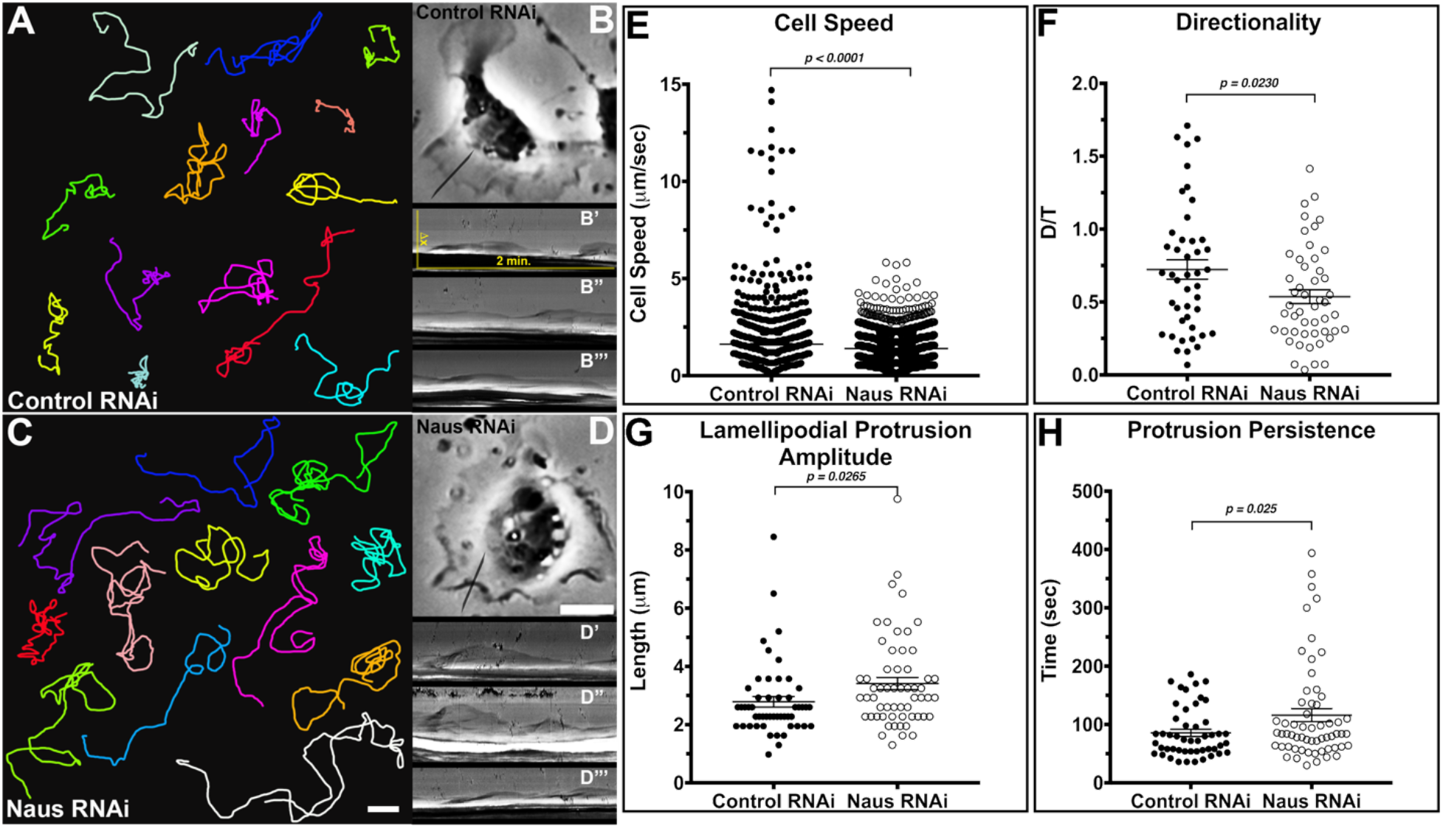
Nausicaa regulates directionality and lamellipodial persistence in migrating D25c2 cells. (A-D) Representative migration tracks for *Drosophila* D25 cells treated with control (A & B) or Naus (C & D) RNAi. (B’-B’’’) Representative kymographs from control RNAi treated D25 cells, the black line represents a typical region of interested used to generate kymographs. (D’-D’’’) Representative kymographs for Naus RNAi treated D25 cells, the black line represents a typical region of interested used to generate kymographs. (E) Average instantaneous cell speeds for control or Naus RNAi treated D25 cells. Naus depletion led to a slight, but statistically significant reduction in cell migration speeds (***p-value <0.0001, Student’s t-test, Control: n = 43 cells, 1438 measurements, Naus RNAi: n = 50 cells, 1779 measurements). (F) Directionality, was measured as the ratio of D/T where D is distance between starting and end point and T is the total distance traveled. Naus depleted cells showed a statistically significant reduction in directionality as compared to control RNAi treated cells (*p-value = 0.0230, Student’s t-test, Control: n = 44 cells, Naus RNAi: n = 49 cells). (G) Naus RNAi led to an increase in the length of lamellipodial protrusions as compared to control RNAi treated D25 cells (*p-value = 0.0265, Student’s t-test, Control: n = 51 cells, Naus RNAi: n = 58 cells). (H) Protrusion persistence was increased upon treatment with Naus RNAi as compared to control treated D25 cells (*p-value = 0.025, Student’s t-test, Control RNAi: n = 49 cells, Naus RNAi: n = 59 cells). All error bars = S.E.M. Scale bar = 10 μm.

While these results suggest that Naus plays a role in maintaining both the speed and the directionality of migrating cells we wanted the further explore how Naus regulates the lamellipodial dynamics that govern cell migration. Using kymographs taken from phase-contrast microscopy movies we measured lamellipodial persistence, the speeds of protrusion and retraction, the frequency of protrusions, and the amplitude of protrusions (Figure 7B & D) (Bear et al., 2002; Hinz et al., 1999). Despite the slower rates of cell migration, Naus depleted cells had longer lamellipodial protrusions that persisted for greater periods of time as compared to control RNAi treated cells. The average maximum length of lamellipodial protrusions (the amplitude of protrusions) for Naus depleted D25 cells was 3.4 ± 0.21 μm which is statistically significantly greater than control cells at 2.8 ± 0.18 μm (p=0.0265, Student’s t-test) (Figure 7G). This data corroborates our ultrastructural data in S2R+ cells and suggests the lamellipodia of Naus depleted D25 cells may also contain longer, unbranched filaments. Interestingly, unlike the less persistent lamellipodia of cells where Ena/VASP proteins were membrane targeted, we observed an increase in persistence following Naus depletion, from 85.8 ± 6.0 s in control RNAi treated cells to 116 ± 11.0 s in Naus RNAi treated cells (p=0.025, Student’s t-test) (Figure 7H). This suggests that despite have long, unbranched filaments, the lamellipodia of Naus depleted cells are still able to protrude without an increase in buckling. Depletion of Naus had a specific effect on lamellipodial dynamics and other parameters such as the frequency of protrusions, the speed of protrusions and retractions, and the total distance the cells migrated over a two-hour period remain unchanged as compared to control RNAi treated D25 cells (Supplemental Figure 4). Collectively, we found that Naus depletion in D25 cells led to an increase lamellipodial persistence while simultaneously decreasing cell migration speeds and directionality.

### Depletion of Nausicaa decreases the number of branches in Drosophila larval neurons

This fine-tuning of actin dynamics is not only critical to the function of lamellipodia, but plays a major role in the morphology and function of other dynamic actin structures. One actin based-structure that is particularly sensitive to changes in actin dynamics are dendritic spines (Fischer et al., 1998; Hotulainen and Hoogenraad, 2010; Matus, 2000). Accordingly, we sought to determine whether Naus also plays a role in morphology of *Drosophila* neurons. While it remains controversial whether *Drosophila* neurons form dendritic spines *in vitro* we focused on the overall neuronal morphology of third instar larvae neurons in culture. Using the UAS-Gal4 system, we depleted Naus specifically in neurons using the pan-neuronal Gal4 driver Elav. The brains of third instar larvae were removed and enzymatically dissociated. The resulting neuroblasts were allowed to differentiate in culture for 24 hours (Lu et al., 2013) (Figure 8A-C).

**Figure 8.**
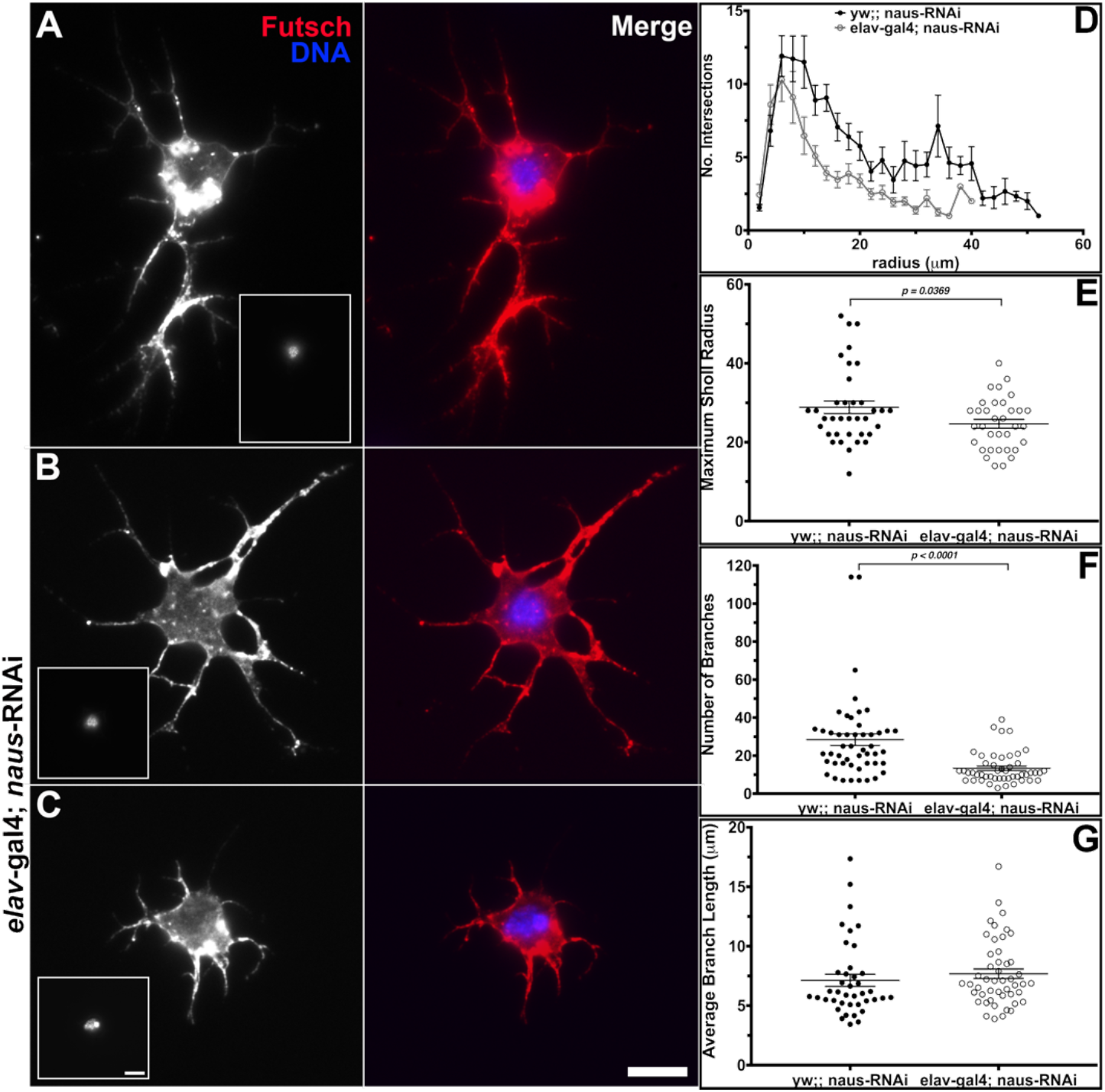
Nausicaa regulates neuronal morphology in 3rd instar larvae neurons. (A-C) Primary neuroblasts from 3rd instar larvae were harvested and allowed to differentiate for 24 hours in culture. They were then fixed and stained for microtubules (anti-Futsch, red in merged images) and DNA (Hoechst, inset and in blue in merged images). (A) Control neurons from yw;; naus-RNAi flies. (B & C) Neurons from flies expressing naus-RNAi driven by elav-gal4. Scale bars = 10 μm.(D) Naus depletion also led to a decrease in the Sholl profile, which measures the number of intersections a neuron makes, (error bars denote S.E.M.). (E) The maximum Sholl radius, which measures the number of branch intersections from concentric circles also indicated a decrease in Naus RNAi neurons (open gray circles) as compared to control neurons (black circles) (*p-value = 0.0369, Student’s t-test, control: n = 36 cells, Naus RNAi: n = 34 cells). (F & G) 2D skeletons of neurons were manually drawn and analyzed ImageJ Simple Neurite Tracer for (F) number of branches and (G) the average branch length. (F) There was a statistically significant decrease in the average number of branches in Naus depleted neurons (open gray circles) as compared to control neurons (black circles) prepared in parallel (***p-value <0.0001, Student’s t-test, Control: n = 49 cells, Naus RNAi: n = 48 cells). (G) The average branch length was not significantly different between control (black circles) and Naus RNAi (open gray circles) neurons. Error bars = S.E.M.

Following fixation and staining with the neuronal marker Futsch, the morphology of the neurons was assessed using Sholl analysis (Ferreira et al., 2014; Sholl, 1953). Sholl analysis analyzes the neuronal morphology by counting the number of intersections for concentric circles from the center of the cell body. Interestingly, we observed a distinct difference in the Sholl profiles of Naus depleted neurons as compared to control neurons prepared in parallel, suggesting a difference in neuronal arborization (Figure 8D). Similarly, the maximum Sholl radius in which neuronal intersections were still detected was significantly lower in Naus depleted neurons (Figure 8E). Consistent with this result, when neuron 2D skeletons were analyzed with ImageJ Simple Neurite Tracer (Longair et al., 2011), we observed that Naus depleted neurons showed decreased average branch length in comparison to controls (Figure 8F & G). Taken together, these results suggest that Naus plays a role in neuronal branch arborization and branch length.

## Discussion

The formation of actin protrusive structures such as the lamellipodia of migrating cells and the dendritic spines of neurons rely on Arp2/3 generated actin branches. Changes to the density and stability of actin branches can affect the overall morphology of these structures and ultimately, their function. We are interested in proteins that play a role in this fine-tuning of actin branches such as the type II NPF and actin branch stabilizer, Cortactin. Here we characterize Nausicaa, a putative *Drosophila* homolog of two mammalian Cortactin binding proteins, CTTNBP2 and CTTNBP2NL. Using cultured and primary *Drosophila* cells we demonstrate that Naus, through its interaction with Cortactin, regulates actin-branch dynamics, lamellipodial protrusion, and the morphology of neurons.

### Nausicca is likely the fly homolog of both CTTNBP2 and CTTNBP2NL

We described a previously uncharacterized protein encoding gene *cg10915*, which we have subsequently named *nausicaa* (*naus*). While bioinformatic queries indicate the *naus* is more closely related to mammalian CTTNBP2NL than CTTNBP2, based on its localization and putative role in regulating the morphology of neurons, we argue that Naus covers the function for both proteins in flies. We do not have to look much further for another example of this refinement of the *Drosophila* genome then the Ena/VASP family of proteins. While there are three of these critical regulators of the actin cytoskeleton in a typical mammalian genome, mammalian ena or Mena, VASP, and EVL (Ena/VASP-like), the fly genome only contains one, Enabled (Ena). It is interesting to speculate that the mammalian homologs could be the result of a gene duplication event of an ancestral gene that is similar to *naus*.

### Nausicca’s lamellipodial enrichment is Cortactin dependent

We used TIRF microscopy of both fixed and live cells in combination with RNAi treatments and point mutations and determined that Naus’ lamellipodial enrichment, is Cortactin dependent. Using two different RNAi sequences targeting Cortactin, we found that depletion of Cortactin leads to a loss of Naus localization at actin structures. This differs from the mammalian counterpart in which re-localization of Cortactin by glutamate stimulation in neurons does not lead to CTTNBP2 redistribution. We observed a similar loss in localization upon expression of a point mutant where the proline residues in the conserved proline patch (Supplemental Figure 1) were mutated to alanine (amino acid positions 563-567). Thus Naus, like its mammalian counterparts, uses this same conserved region for its association with Cortactin. Along with abolishing the interaction with Cortactin in COS cells, a similar proline mutant of rat CTTNBP2 failed to rescue the decrease in dendritic spine density following depletion of endogenous CTTNBP2 (Chen et al., 2012), arguing that not only is this association important for proper localization, but it is also needed for proper function.

### Nausicaa and Cortactin regulate one another’s dynamics in a reciprocal manner

Our kinetic studies of Naus and Cortactin reveal a mutual relationship between the two proteins. By both FRAP and Permeabilization Activated Reduction in Fluorescence (PARF), we found that following Cortactin RNAi, Naus was no longer anchored to the actin cytoskeleton. This kinetic data supports our localization studies and further implicates Cortactin’s role in recruiting Naus to the cytoskeleton. Cortactin is incorporated throughout the lamellipodia where it likely targets nascent actin branch junctions having a 300-fold increased affinity for junctions over the sides actin filaments. Interestingly, using *in vitro* single molecule imaging, Helgeson and Nolen observed that Cortactin’s average lifetime at existing branch junctions during active branching was 29.5 s (Helgeson and Nolen, 2013), while the lifetime of branches *in vitro* has been observed to be between 8 and 27 minutes (Mahaffy and Pollard, 2006; Martin et al., 2006). Our kinetic data (Figure 4) suggests that Naus may function to retain Cortactin throughout the lamellipodia, likely at branch junctions given Cortactin’s high affinity for these sites. While this increase in the time at junctions may not be on par with the reported lifetimes *in vitro*, lamellipodial actin undergoes treadmilling. Thus, this delay in the dissociation of Cortactin as a result of its association with Naus may be more consequential *in vivo*. Delays in Cortactin’s dissociation could also lead to a decrease in the number of new branches given its ability to function as a type II NPF. Furthermore, while outside the scope of this study, it would be interesting to determine how Naus affects the activity of de-branching enzymes such as cofilin or GMF given Naus’ putative role in regulating branch formation and dynamics.

### Naussica regulates branch density, actin-retrograde flow rates, and lamellipodial protrusion

Actin retrograde flow is the result of both contractility generated by non-muscle myosin II and the force of the cell membrane pushing back against actin filaments as they polymerize. Quantitative fluorescence speckle microscopy revealed that depletion of Naus leads to a decrease in actin retrograde flow speeds within the lamellipodia (Figure 3). This decrease in retrograde flow may be the result of an overall decrease in Arp2/3 nucleation and branch junction stabilization. Two critical observations led us to this hypothesis. Firstly, our kinetic data indicates Cortactin more readily dissociates from the actin cytoskeleton following Naus depletion. Secondly, we found that RNAi depletion of Naus led to an increase in actin in the number of long, unbranched actin filaments by platinum replica electron microscopy (Figure 6). Thus, Cortactin could be undergoing a cycle of precocious dissociation from branch junctions leading to less activation of the Arp2/3 complex and a decrease in the stability of branched junctions. This putative mechanism draws parallels to what has been observed in fibroblasts lacking Ena/VASP proteins (Bear et al., 2002). The lamellipodia of fibroblast where Ena/VASP proteins were miss-targeted to the cell membrane and contained longer, less branched actin filaments. These changes to the actin architecture resulted also affected the dynamics of the lamellipodia. The longer, less branched actin filaments decreased the speed of cell migration and the directionality of this migration but increased the amplitude of lamellipodial protrusions and increased lamellipodial persistence (Figure 7). Interestingly, depletion of Cortactin also led to a decrease in random cell motility, however these cells had less persistent lamellipodial protrusions suggesting nuanced differences between the loss of Cortactin and the loss of Naus to actin dynamics (Bryce et al., 2005). These differences may very well lie in function of Naus to stabilize Cortactin retaining it in the lamellipodia. Naus depletion appears to differ somewhat from the depletion of the branch destabilizer Coronin 1B as well. The depletion of Coronin 1B leads to a more densely branched actin network and a decrease in retrograde flow rates. Coronin 1B depletion also leads to an increase the speeds of lamellipodial protrusion while reducing lamellipodial persistence (Cai et al., 2007; Hostos et al., 1993; Krause and Gautreau, 2014). Thus, Naus’ role in fine tuning the lamellipodia is distinct from that of both Cortactin and Coronin 1B.

### Naussica plays a role in regulating neuronal morphology

This role in regulating actin dynamics also plays out in determining the morphology of neurons. Using fly genetics we depleted Naus in neuroblasts, which differentiate into neurons in culture. We found that depletion of Naus led to a decrease in the number of processes made in comparison to wild-type neurons (Figure 8). Similarly, depletion of CTTNBP2 also led to a decrease in neuronal arborization as well as a decrease in the density of dendritic spines (Chen and Hsueh, 2012; Chen et al., 2012; Shih et al., 2014). However, CTTNBP2 also promotes microtubule stability, thus its role in promoting neuronal arborization may have diverged from its role in regulating Cortactin dynamics (Shih et al., 2014). Interestingly, we did not observe wild-type Naus associating with microtubules and it was only upon expression of the alanine mutant (Naus-AAAIA, Supplemental Figure 2D), albeit on a rare occasion, did we observe colocalization with microtubules. Understanding the differences between Naus and CTTNBP2 will likely be the focus of future studies, in particular if they both contribute to the morphology of neurons in distinct ways despite being closely related.

### Working model for Nausicaa’s role in branch nucleation, stabilization and the fine-tuning of the lamellipodia

Given the observations detailed here, we propose a model wherein Nausicaa acts through the stabilization of Cortactin at Arp2/3 generated branches to regulate their dynamics (Figure 9). By stabilizing Cortactin, Naus inhibits its precocious dissociation while preventing debranching. Without Naus, Cortactin more freely dissociates from Arp2/3 generated branches leading to the loss of destabilization of branch junctions and an overall decrease in actin branch density throughout the lamellipodia. This decreased density leads to larger scale cellular changes, such as reduced speeds in cell migration (Figure 7) and decreased neuronal branches observed in this study (Figure 8). In a similar manner, Cortactin maintains Naus at the lamellipodia, and when Cortactin is absent, Naus loses this enrichment and, in extremely rare cases, relocalizes to other structures such as microtubules. Collectively, both Naus and Cortactin act in concert to ensure the appropriate spatial and temporal regulation of lamellipodial actin dynamics.

**Figure 9.**
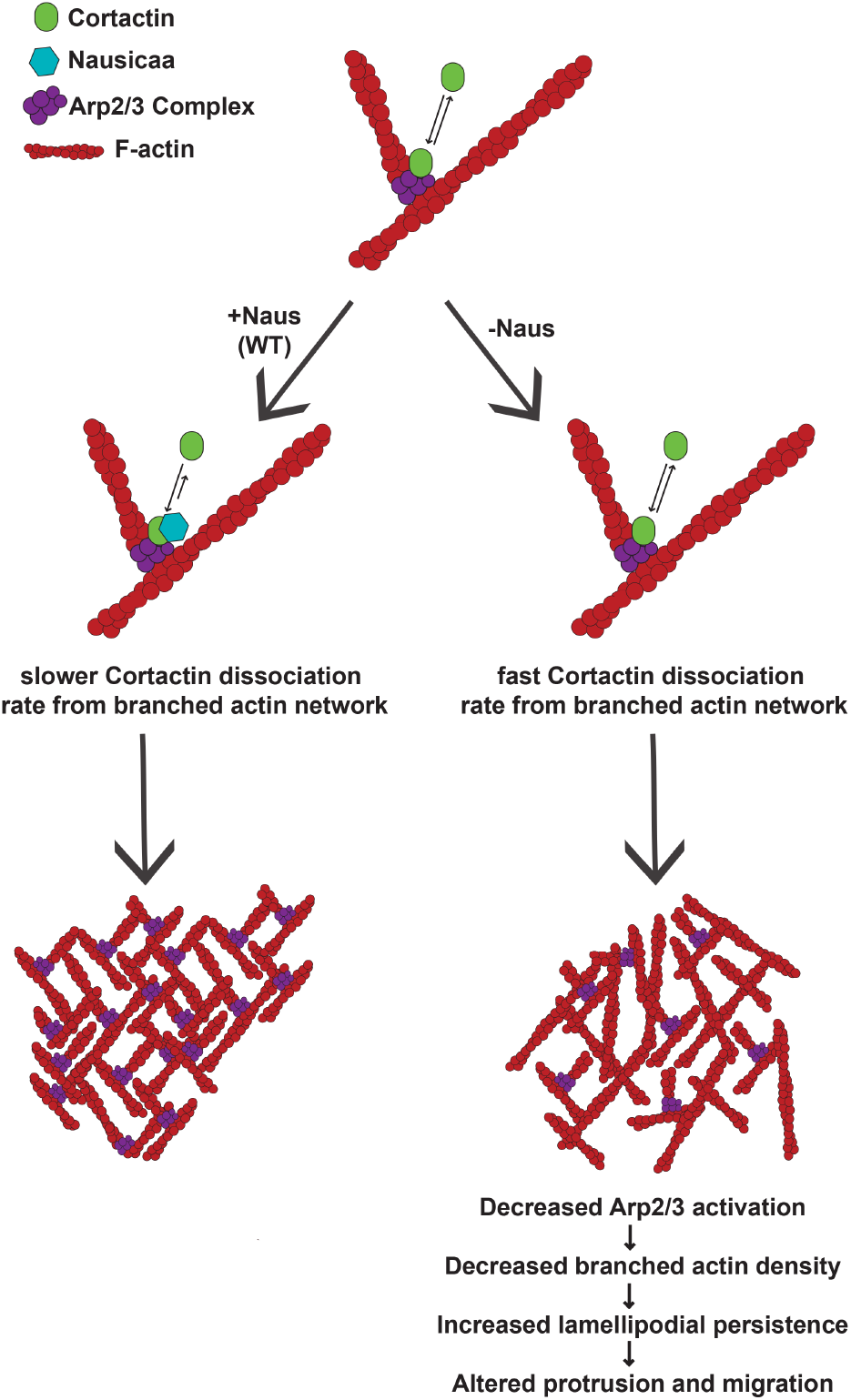
Proposed model of Nausicaa’s role in the lamellipodia. Nausicaa works to stabilize Cortactin at Arp2/3 generated branches in order to appropriately regulate branch density of the lamellipodia. Nausicaa’s interaction with Cortactin maintains Cortactin on branched actin networks and stabilizes these junctions. In the absence of Nausicaa, Cortactin has a fast off-rate from Arp2/3 branches (Helgeson and Nolen, 2013). Cortactin more freely diffuses leading to a decrease in the activation of the Arp2/3 complex and a loss of branch stability. This increased filament length eads to larger lamellipodial protrusions and downstream alterations to migration and morphology.

## Materials and Methods

### Cell Culture and RNAi

Drosophila S2R+ cell culture and RNAi were performed as described in Rogers and Rogers, 2008 and Applewhite et al., 2016. Briefly, S2R+ (Drosophila Genomics Resource Center, Bloomington, IN) cells were cultured in Shields and Sanger media (Sigma-Aldrich, St. Louis, MO) supplemented with 100X antibiotic-antimycotic (Thermo Fisher Scientific, Waltham, MA), and 10% fetal bovine serum (Thermo Fisher Scientific, Waltham, MA) maintained at 25°C.

RNAi was administered in six-well plates by treating cells (approximately 50% confluent) with 10 μg of double-stranded RNA (dsRNA) in 1 ml of medium each day for 7 days. Control RNAi was made from dsDNA amplified from pBlueScript vector with no known homology to the *Drosophila* genome. For all other dsRNA targets please see Supplemental Table 1 for primer sequences.

*Drosophila* ML-DmD25c2 (D25 cells, Drosophila Genomics Resource Center, Bloomington, IN) were maintained as described in Currie and Rogers, 2011. Briefly, D25 cells were cultured in Schneider’s media (Thermo Fisher Scientific, Waltham, MA) supplemented with 100X antibiotic-antimycotic (Thermo Fisher Scientific, Waltham, MA), 10% fetal bovine serum (FBS, Thermo Fisher Scientific, Waltham, MA), and 10μg/ml insulin (Thermo Fisher Scientific, Waltham, MA). RNAi regimen was the same as described for S2R+ cells (see above).

*Drosophila* primary neuroblasts were harvested and cultured as described in Lu et al., 2013. Briefly, the brains of third instar larvae were dissected in Schneider’s media supplemented with 20% FBS, and then enzymatically dissociated with liberase (Roche, Basel, Switzerland) at a final concentration of 0.20-0.25 mg/ml in Modified Dissecting Saline (137 mM NaCl, 5.4 mM KCl, 0.17 NaH2PO4 0.22 mM HKPO4 3.3 mM Glucose, 43.8 mM Sucrose, 9.9 mM Hepes, pH 7.5). The Modified Dissecting Solution was replaced with Schneider’s media supplemented with 20% FBS and neuroblasts were plated on ECM harvested from the D25 cells (see Currie and Rogers, 2011) and allowed to differentiate for 24 hours at 25°C.

### Molecular Biology

The cDNA clones for Nausicaa (CG10915) and Cortactin (CG3637) were obtained from the Drosophila Genomics Resource Center (University of Indiana, Bloomington, IN) and were cloned into pMT or pIZ (Invitrogen) vectors following standard PCR procedures. Nausicaa’s conserved Cortactin binding motif (amino acid positions 563-567) were mutated to alanine by site directed mutagenesis.

### Immunofluorescence and Live-cell Imaging

Cells were prepared for immunofluorescence and live-cell imaging as described in Applewhite et al., 2016. S2R+ cells were plated on concanavalin A-treated coverslips attached to laser cut 35 mm-tissue culture dishes with UV-curable adhesive (Norland Products, Cranbury, NJ) in Shields and Sanger media supplemented with 10% FBS and 100X antibiotic-antimycotic for both fixed and live-cell imaging. D25 cells were plated on glass bottom dishes (described above) treated with ECM harvested from the cells as described in Currie and Rogers, 2011. Antibodies used in this study include anti-Myc 9E10 (Developmental Hybridoma Bank, Iowa City, Iowa), anti-Futsch (Developmental Hybridoma Bank, Iowa City, Iowa), anti-alpha tubulin (Developmental Hybridoma Bank, Iowa City, Iowa), and anti-beta tubulin (Developmental Hybridoma Bank, Iowa City, Iowa) diluted 1:200 in a 5% solution of normal goat serum (Sigma-Aldrich) and phosphate-buffered solution with 0.1% Triton x-100 (PBST) (Sigma-Aldrich). Secondary antibodies (Alexa-488 and Alexa 594; Jackson ImmunoResearch, West Grove, PA) and phalloidin (Alexa-488 and Alexa-594; Thermo Fisher Scientific) were used at final dilution of 1:100 in PBST. Hoechst (Thermo Fisher Scientific) was diluted 1:10,000 in PBST. All transfections were carried out using using FuGENE HD (Promega, Madison, WI). Expression of pMT vectors was achieved with 250-500 μM final concentration of copper sulfate unless noted otherwise. Cells were fixed using a 10% solution of Paraformaldehyde (Electron Microscopy Sciences, Hatfield, PA) and PEM buffer (100 mM Pipes, 1 mM EGTA, 1 mM MgCl_2_). Fixed cells were mounted using Dako anti-fade mounting media (Agilent, Santa Clara, CA). All imaging was performed on a total internal reflection fluorescence (TIRF) system mounted on an inverted microscope (Ti-E, Nikon, Tokyo, Japan) using a 100X/1.49NA oil immersion TIRF objective driven by Nikon Elements software unless noted otherwise. Images were captured using a Orca-Flash 4.0 (Hamamatsu, Hamamatsu, Japan) and were processed for brightness and contrast using ImageJ before analysis.

### Co-localization Analysis

Co-localization was analyzed by line-scan analysis and Mander’s coefficient analysis. For line-scan analysis, a 10 μm line was drawn from the cell edge inward and fluorescence intensity was measured. These values were normalized and then averaged for all cells within that condition. Mander’s coefficient analysis was performed using the Just Another Colocalization Program (JACoP) plugin for ImageJ (Bolte and Cordelières, 2006). Briefly, intensity thresholds were manually set for both fluorescence channels and then the fraction of overlap was calculated in each direction.

### Neuroblast Analysis

Neuroblasts were analyzed using the Simple Neurite Tracer and Sholl Analysis plugins in ImageJ (Ferreira et al., 2014; Longair et al., 2011). For Sholl Analysis, neuroblasts were converted to a threshold image. Following, a line from the center of the soma to past the further branch tip was drawn to define the space for analysis. The radius for analysis was set to 2 μm concentric circles and the number of intersections per radius was calculated. Similarly, the max Sholl radius was then extracted by the maximum radius at which the number of intersections was greater than zero. For analysis of average branch length and number of branches using Simple Neurite Tracer, a line skeleton of the neuron image was manually drawn and these values were then calculated using the plugin.

### Permeabilization Activated Reduction in Fluorescence (PARF)

PARF was performed as described in Singh et al., 2016. Briefly, cells were prepared for live imaging as described above. Time-lapse images were captured with constant exposure at a rate of one frame every two seconds. After 40 seconds (20 frames), digitonin (25 μM final concentration) was added. Cells from the same dish were imaged under the same conditions but without digitonin treatment for use as a photofading due to acquisition (PDA) control. Analysis was performed using ImageJ and GraphPad Prism 6. The area of each region of interest (ROI) was held constant. An ellipse of the background of each movie (control and digitonin treatment) was selected and intensity density was determined for the background in each frame. Photofading due to acquisition (PDA) was determined as previously described in Applewhite et al., 2007. The intensity density was determined for a lamellipodial ROI in the control (no digitonin treatment) movie for each plate. The background intensity was subtracted and change in fluorescence was fit to a one phase exponential decay of the following general equation where I is intensity and k is the photofading factor: I = e^−kt^ + I_o_. The intensity density of a lamellipodial ROI for digitonin cells was obtained in the same manner. Background intensity was subtracted and the intensity density was then multiplied by e^kt^. The intensity was normalized for each cell and the data was averaged for each condition. To compare the half-life between conditions for statistical significance, the normalized fluorescence for each cell was fit to a one phase exponential decay and t_1/2_ was determined. These half-life values were averaged in each condition and compared using a Student’s t-test.

### Fluorescence Recovery after Photobleaching (FRAP)

FRAP was performed using a Zeiss LSM880 laser-scanning confocal microscope (Jena, Germany). Cells were prepared for live imaging as described above. Time-lapse images were captured every 1.34 seconds. After 50 cycles (65.6 sec), selected regions of the cell were bleached (5 iterations) and the intensity was recorded. Intensity was also recorded for a nonbleached region and a background region of the same size was uses for PDA controls. FRAP analysis was performed as described in Applewhite et al., 2007. The fluorescence intensities in the bleached zone in each frame were measured. The background was subtracted and the intensity was corrected for photofading as described above. The intensity was normalized for each ROI. This corrected intensity was fit to a one phase association. The half-life of recovery was calculated as ln2/k. Values were compared using a Student’s t-test.

### Quantitative Fluorescence Speckled Microscopy (QFSM)

RNAi treatment and transfection was performed as described above. Following transfection, cells were induced with 30μM copper sulfate and incubated overnight. Live cell movies were obtained at 200 ms exposure in 2 sec intervals for 2 min. The resulting movies were analyzed using a previously described Quantitative Fluorescent Speckle Microscopy (QFSM) software in MATLAB (Mendoza et al., 2012). Images were acquired at a rate of 30 frames per minute (130nm/pixel, NA=1.4, 16 bit images). Full cell masks were generated using automatic thresholding (MinMax setting). Flow analysis was performed using the flow tracking setting with 6 frame integration window. For cell-wise quantification of lamellipodial flow rates, masks of lamellipodial regions for each cell were generated and the average actin flow rate was calculated for the first 30 seconds of each video (n=27-35 cells / group).

### Platinum Replica Electron Microscopy

Sample preparation for platinum replica electron microscopy was performed as previously described in Svitkina, T., 2016. Briefly, cells were extracted in Extraction buffer (1% Triton X-100, 2% PEG (MW 35 kDa) in PEM buffer supplemented 2 μM phalloidin), washed with PBS and then fixed with 2% glutaraldehyde (EM grade from Electron Microscopy Sciences, Hatfield, PA) in 0.1 M sodium cacodylate, pH 7.3. Fixed cells were then treated with 0.1% tannic acid and 0.2 % uranyl acetate in water, critical-point dried, and coated with platinum and carbon. They were then transferred to EM grids for imaging.

### Random Cell Migration Assay and Kymography

D25 cells were plated at a subconfluent density on ECM coated glass bottom dishes and allowed to attach overnight. Cells were imaged every 5 minutes for 6 hours by phase-contrast microscopy using 40X/0.75NA objective. Individual cells were manually tracked using Manual Tracker (ImageJ). Cell directionality was calculated as a ratio of the direct distance between start and end points (D) to the total path length taken by the cells (T). To measure the rates of lamellipodial protrusion, retraction, persistence, frequency, and amplitude, kymographs were made using the Multi Kymograph ImageJ plugin from phase-contrast movies acquired every 2 seconds for 10 minutes. Kymographs were generated from phase-contrast movies of migrating D25 cells acquired every 2 seconds for 10 minutes. A line approximately 16 microns in width was drawn from the center of the cell to a few microns beyond the cell periphery. Following the protocol established by Hinz et al., 1999, these kymographs were used to extract the lamellipodial protrusion parameters. Statistics were performed using GraphPad Prism 7.

## Acknowledgements

We would like to acknowledge the help of Dr. Stephanie Kaeche Petrie at Oregon Health and Science University for her mentorship and training on the FRAP experiments, and Dr. Farida Korbova for the electron microscopy and fruitful discussions. In addition we would like to thank Drs. Ryan Fink, Joseph Lee, and J. Clapp for their assistance in transcontinental fly transportation, Drs. Jeremy Coate and Anna Ritz for their assistance with bioinformatics, the Drosophila Genomics Resources Center (NIH grant 2P40OD010949 to DGRC) and Developmental Studies Hybridoma Bank for reagents. Additionally, we would like to thank Mari Cobb, Emily Merfeld, and Abrar Abidi for their input and helpful discussions throughout this project, and Drs. Vladimir Gelfand, Stephen Rogers, and Omar Quintero for careful reading of this manuscript.

## Funding Information

This work was supported by the National Science Foundation STEM Grant (NSF1154004 to Reed College), the Reed College Biology Undergraduate Research Fellowship (to M.E.O.), the Reed College Science Research Fellowship (to M.E.O.), the National Institutes of Health (R15 GM122019-01 to D.A.A), the National Science Foundation (NSF 716964 to D.A.A.) and generous start-up funds from Reed College (to D.A.A.).

## Competing Interests

The authors declare that there are no competing interests.

## Supplemental Figures and Tables

**Supplemental Table 1.**
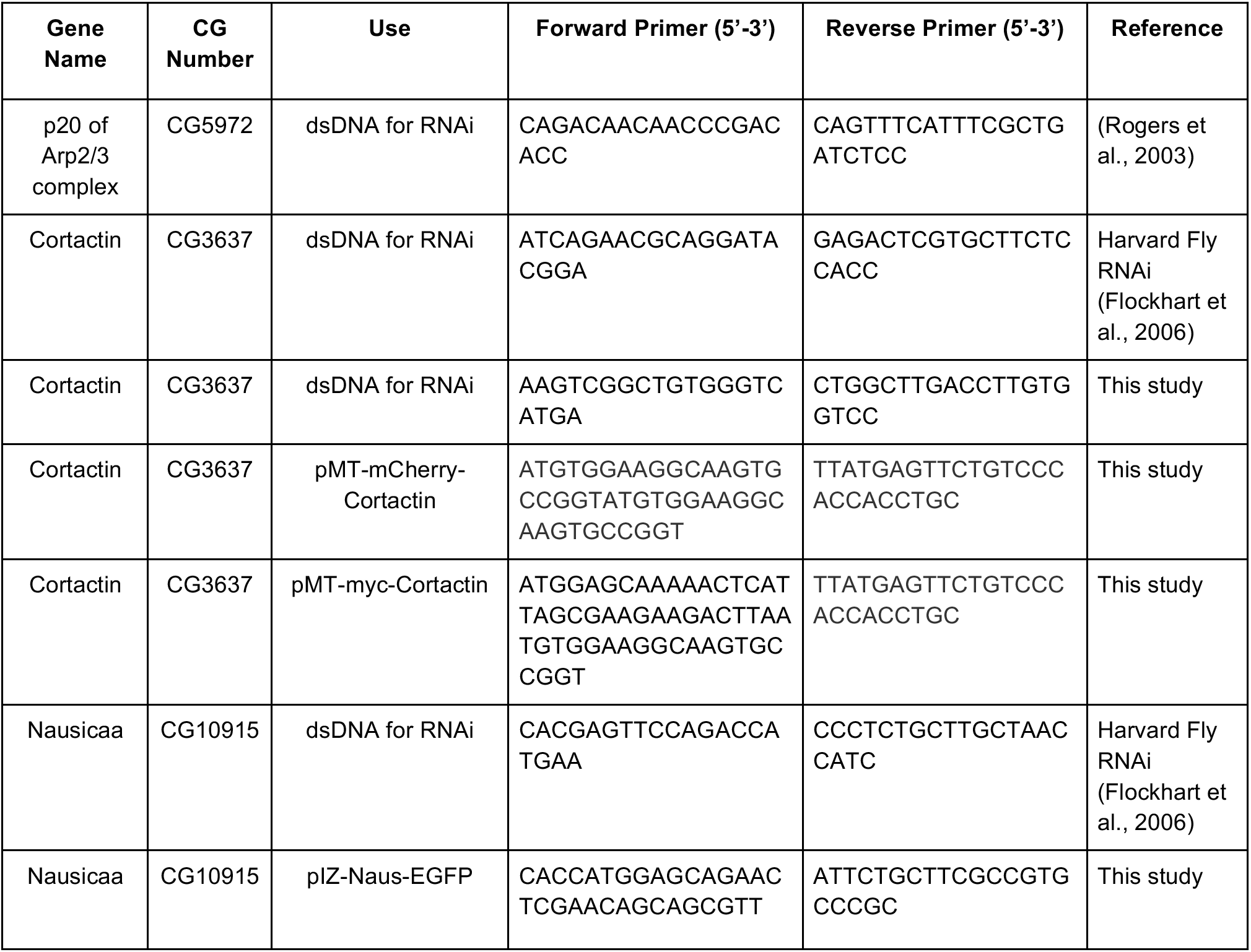
Forward and reverse primer sequences for production of cloning and dsDNA (for dsRNA production) used in this study.

**Supplemental Figure 1.**
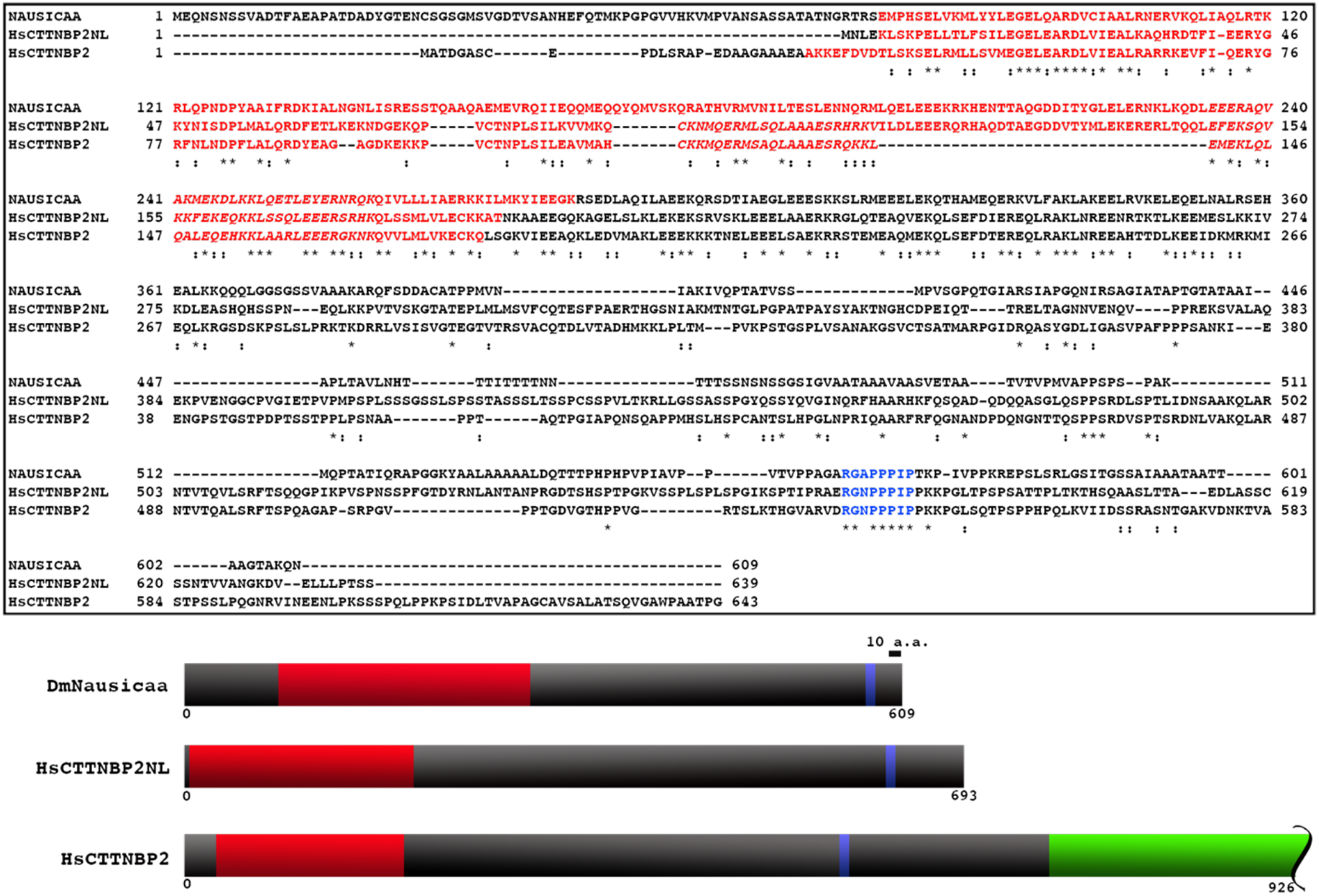
Sequence Alignment of *Drosophila* Nausicaa with *Homo sapien* Cortactin binding protein 2 (CTTNBP2) and Cortactin binding protein 2 N-terminal like (CTTNBP2NL). (Top) A multiple sequence alignment made using Clustal Omega comparing *Drosophila melanogaster (Dm)* Nausicaa (CG10915) and *Homo sapien (Hs)* CTTNBP2NL, and CTTNBP2. Nausicaa shares approximately 30% identity with *Hs* CTTNBP2NL and 28% identity with *Hs* CTTNBP2. Shown in red is the conserved Cortactin Binding Protein-2 (CortBP2) domain, in red italics a predicted a coiled-coil domain, and in blue the poly-proline Cortactin binding motif. Asterisks indicate identical amino acids while colons indicate similar amino acids. Note that only the first 643 amino acids of *H.s*. CTTNBP2 are shown. (Bottom) Diagram of *Dm* Nausicaa, *Hs* CTTNBP2NL and *Hs* CTTNBP2 drawn to scale. In red is the conserved CortBP2 domain, in blue the Cortactin binding motif, and in green the COOH-terminus of CTTNBP2. Note that only the first 926 amino acids of CTTNBP2 are shown.

**Supplemental Figure 2.**
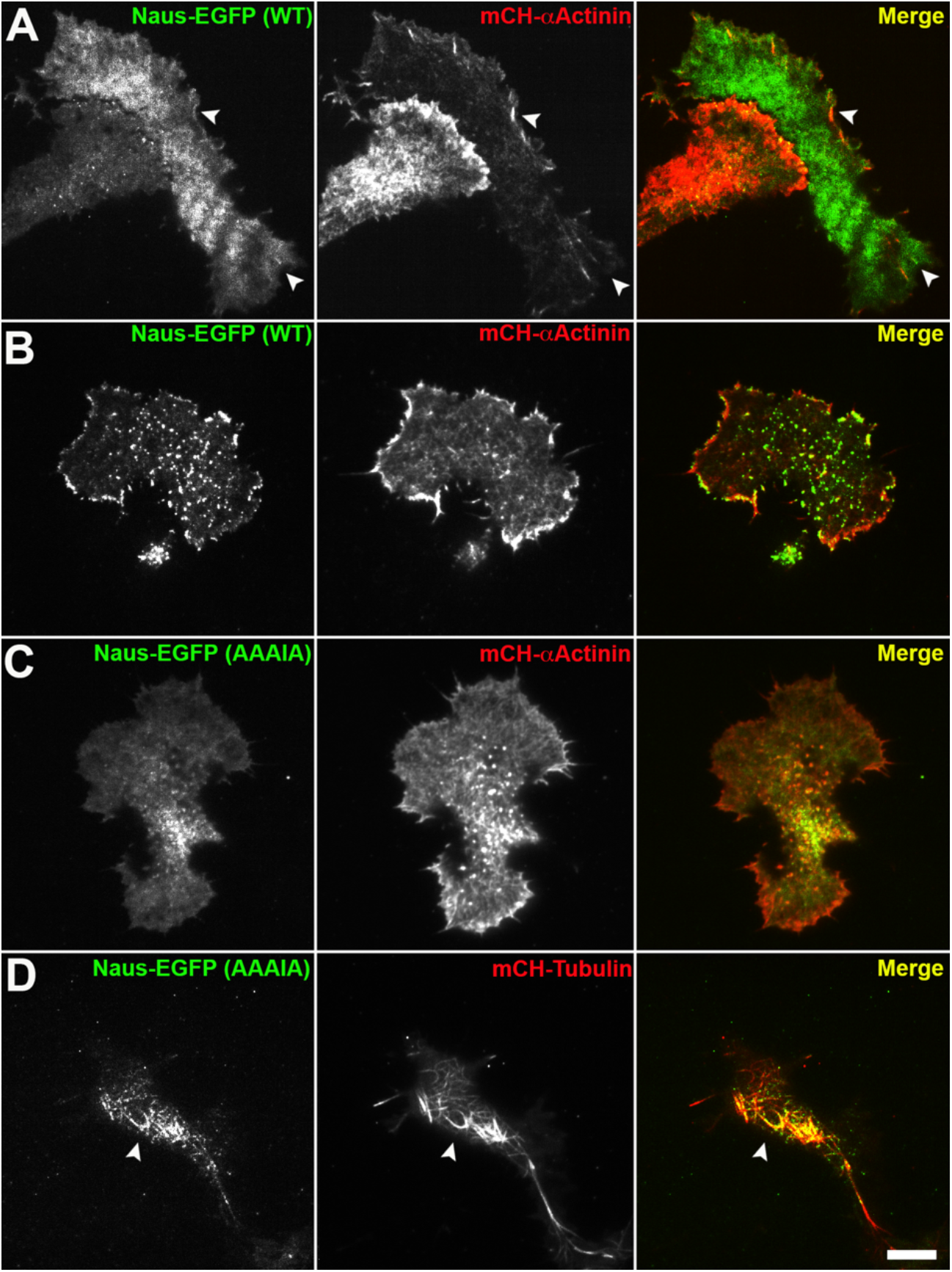
Nausicaa weakly localizes to α-Actinin bundles in D25c2 cells through a proline rich motif, while mutant Naus localizes to microtubules. (A-D) Representative live-cell images of D25 cells cotransfected with mCherry-αActinin (A-C), or mCherry-Tubulin (D) (middle panels, red in merged images) and Naus-EGFP WT (A & B) or Naus AAAIA mutant (C & D) (left panels, green in merged images). (B) White arrowheads indicate bundled-actin structures containing both mCherry-α-Actinin and Naus-EGFP. (C) Note the loss of lamellipodial enrichment in cells expressing Naus-EGFP (AAAIA). (D) White arrowheads indicate where Naus-EGFP (AAAIA) appears to localize to microtubules. Scale bar = 10 μm.

**Supplemental Figure 3.**
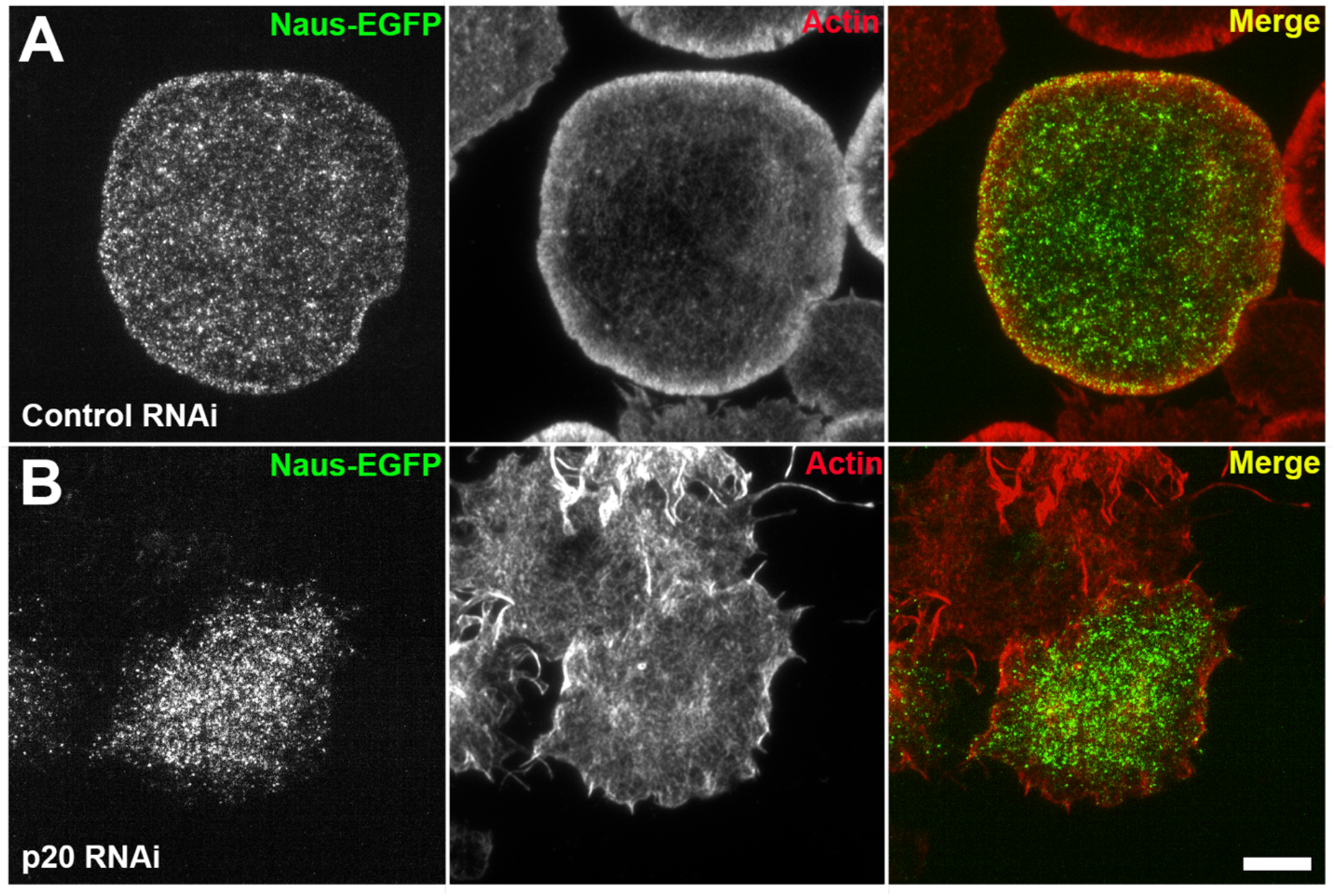
Nausicaa’s lamellipodial localization is dependent on Arp2/3 complex. Fixed S2R+ cells expressing Naus-EGFP (left panels, green in merged images) and treated with either control (A) or p20 (B) RNAi. Cells are stained for F-actin with phalloidin (middle panel, red in merged images). Scale bar = 10 μm.

**Supplemental Figure 4.**
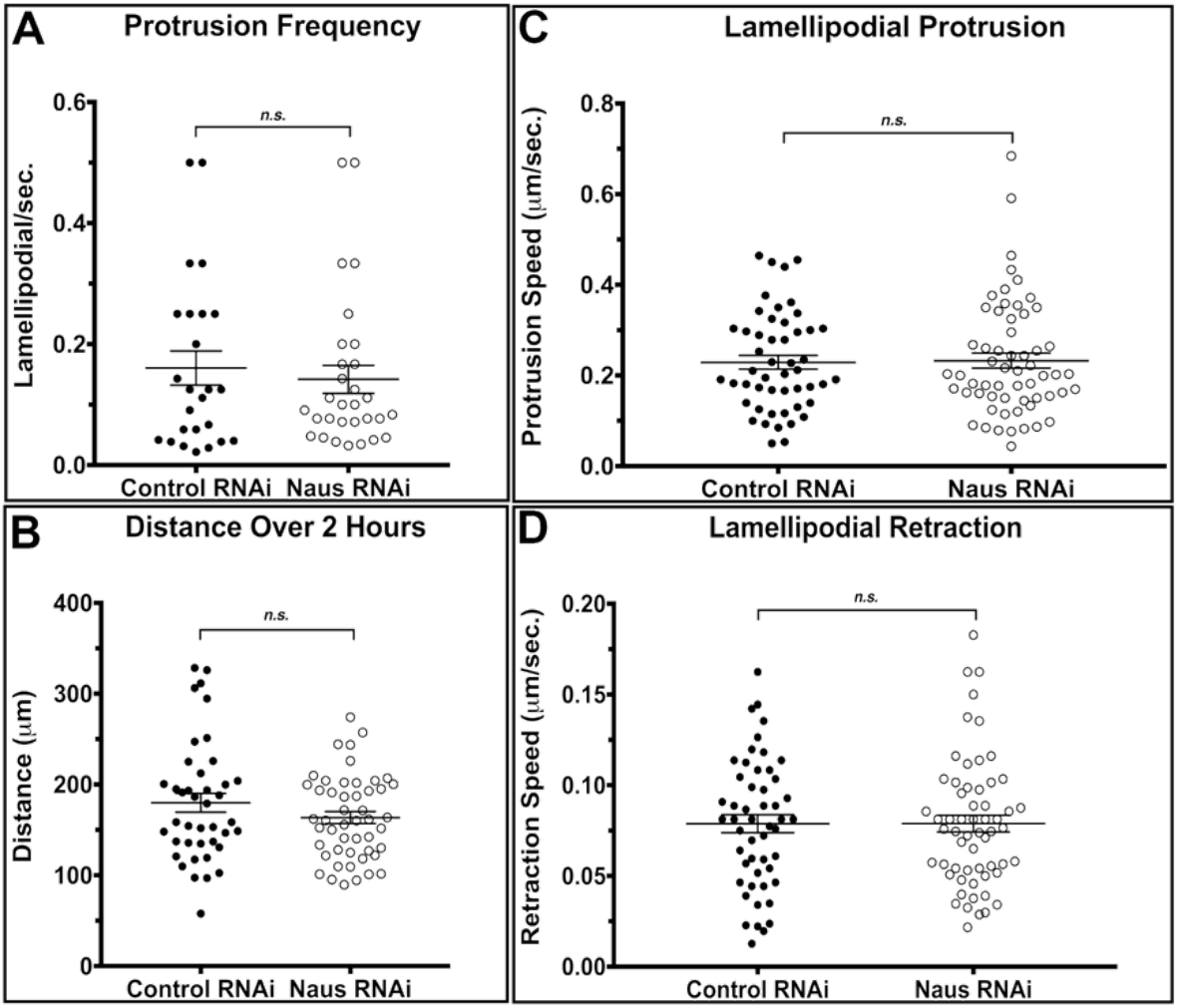
Nausicaa regulates specific lamellipodial parameters. The parameters of lamellipodial dynamics such as frequency of protrusions (A), the distance the cells traveled over two hours (B) the speed of protrusions (C), and the speed of retractions (D) were not statistically significantly different between control and Naus treated D25 cells. N= 51 control cells and 59 Naus RNAi treated cells (A,C, and D) or N= 44 control cells and 50 Naus RNAi treated cells (B).

## Supplemental Videos

All time-lapsed images are played at a rate of 7 frames per second. All images were acquired by TIFR microscopy unless otherwise noted.

**Video 1. Lamellipodial localization of Naus is Cortactin-dependent.** S2R+ cells expressing EGFP-tagged Naus following control RNAi (right) or Cortactin RNAi (left). Image sequence was acquired at 3 second intervals.

**Video 2. PARF of Naus following treatment with control RNAi.** An S2R+ cell expressing EGFP-tagged Naus following control RNAi. After 40 seconds of imaging, the cell was permeabilized with 25 μM digitonin. Image sequence was acquired at 2 second intervals.

**Video 3. PARF of Naus following treatment with Cortactin RNAi.** An S2R+ cell expressing EGFP-tagged Naus following Cortactin RNAi. After 40 seconds of imaging the cell was permeabilized with 25 μM digitonin. Image sequence was acquired at 2 second intervals.

**Video 4. FRAP of Naus following treatment with control RNAi.** An S2R+ cell expressing EGFP-tagged Naus following control RNAi. The cell was photobleached in the regions denoted by the white boxes. The cell was imaged by an LSM 880 confocal microscope at 2 second intervals.

**Video 5. FRAP of Naus following treatment with Cortactin RNAi.** An S2R+ cell expressing EGFP-tagged Naus following Cortactin RNAi. The cell was photobleached in the regions denoted by the white boxes. The cell was imaged by an LSM 880 confocal microscope at 2 second intervals.

**Video 6. PARF of Cortactin following treatment with control RNAi.** An S2R+ cell expressing mCherry-Cortactin following treatment with control RNAi. After 40 seconds of imaging the cell was permeabilized with 25 μM digitonin. Image sequence was acquired at 2 second intervals.

**Video 7. PARF of Cortactin following treatment with Naus RNAi.** An S2R+ cell expressing mCherry-Cortactin following treatment with Naus RNAi. After 40 seconds of imaging the cell was permeabilized with 25 μM digitonin. Image sequence was acquired at 2 second intervals.

**Video 8. QFSM of EGFP-actin.** Using a metallothionein promoter we titrated the addition of copper sulfate in order to generate actin speckles in S2R+ cells. When then imaged at 2 second the resulting actin dynamics following control (left) and Naus RNAi (right). Image analysis was carried out in Matlab.

**Video 9. Random cell motility assay.** D25 cells were treated with control or Naus RNAi for seven days and then were imaged by phase-contrast microscopy over a period of six hours. Image sequence was acquired at 5 minute intervals.

